# Human footprint and forest disturbance reduce space use of brown bears (*Ursus arctos*) across Europe

**DOI:** 10.1101/2024.08.14.607538

**Authors:** Anne G. Hertel, Aida Parres, Shane C. Frank, Julien Renaud, Nuria Selva, Andreas Zedrosser, Niko Balkenhol, Luigi Maiorano, Ancuta Fedorca, Trishna Dutta, Neda Bogdanović, Natalia Bragalanti, Silviu Chiriac, Duško Ćirović, Paolo Ciucci, Csaba Domokos, Mihai Fedorca, Stefano Filacorda, Slavomir Finďo, Claudio Groff, Miguel de Gabriel Hernando, Djuro Huber, Georgeta Ionescu, Klemen Jerina, Alexandros A. Karamanlidis, Jonas Kindberg, Ilpo Kojola, Yorgos Mertzanis, Santiago Palazon, Mihai I. Pop, Maria Psaralexi, Pierre Yves Quenette, Agnieszka Sergiel, Michaela Skuban, Diana Zlatanova, Tomasz Zwijacz-Kozica, Marta De Barba

## Abstract

Three-quarters of the planet’s land surface has been altered by humans, with consequences for animal ecology, movements and related ecosystem functioning. Species often occupy wide geographical ranges with contrasting human disturbance and environmental conditions, yet, limited data availability across species’ ranges has constrained our understanding of how human impact and resource availability jointly shape intraspecific variation of animal space use. Leveraging a unique dataset of 752 annual GPS movement trajectories from 370 brown bears (*Ursus arctos*) across the species’ range in Europe, we investigated the effects of human impact (i.e., human footprint index), resource availability, forest cover and disturbance, and area-based conservation measures on brown bear space use. We quantified space use at different spatio-temporal scales during the growing season (May - September): home range size; representing general space requirements, 10-day long-distance displacement distances, and routine 1-day displacement distances. We found large intraspecific variation in brown bear space use across all scales, which was profoundly affected by human footprint index, vegetation productivity, and recent forest disturbances creating opportunity for resource pulses. Bears occupied smaller home ranges and moved less in more anthropized landscapes and in areas of higher resource availability. Forest disturbances reduced space use while contiguous forest cover promoted longer daily movements. The amount of strictly protected and roadless areas within bear home ranges were too small to affect space use. Anthropized landscapes may hinder the expansion of small and isolated populations, such as the Apennine and Pyrenean, and obstruct population connectivity, for example between the Alpine or Carpathian with the Dinaric Pindos populations. Our findings call for actions to maintain bear movements across landscapes with high human footprint, for example by maintaining forest integrity, to support viable bear populations and their ecosystem functions.

## 1. Introduction

Anthropogenic effects, including climate change, land conversion, fragmentation, and human disturbance are affecting all aspects of animal ecology (Abrahms et al., 2023; Gaynor et al., 2018; Humphries et al., 2002; Prugh et al., 2008). Movement is an integral part of animal ecology and evolution, influencing individuals’ survival and reproduction and more generally, ecosystem interactions, population connectivity and species’ geographic distribution (Nathan et al., 2008; Nathan et al., 2022). Movement facilitates range shifts and allows animals to adapt to climate and global changes (Ellis-Soto et al., 2023; Kauffman et al., 2021). For the last decades, understanding why space use varies within species has received considerable attention (Shaw, 2020). Yet, due to the lack of movement data availability across species’ ranges, the majority of studies have mostly focused on single populations or limited geographic extents (Saïd and Servanty, 2005), restricting our understanding of intraspecific variation in animal movement (Kays et al., 2015). To address this issue, comparative studies integrating large-scale spatial data have contributed toward gaining an understanding of the ecological drivers of animal movements at continental and global scales (Morellet et al., 2013; Mumme et al., 2023; Tucker et al., 2018; Tucker et al., 2023).

How animals use space is largely determined by their motivation and capabilities to move, paired with the prevailing environmental conditions they are exposed to (Morellet et al., 2013; van Beest et al., 2011). Within a species, state-dependent variables such as body size, sex, age, or reproductive status affect energetic needs and the motivation to move (Nilsen et al., 2005a; Tucker et al., 2014). However, because many species have wide geographic distributions that span over contrasting environmental conditions, populations of a given species may exhibit different patterns of space use, with resource availability and seasonality being the most commonly reported factors underlying intraspecific space use variation (Mcloughlin et al., 2000; Olsson et al., 2006; Teitelbaum et al., 2015). In general, where resources are more predictable and abundant, animals tend to occupy smaller ranges, because they can satisfy their energetic requirements within a smaller area (Morellet et al., 2013). Additionally, climatic variables, such as temperature and snow cover and terrain topography also influence animal space use (Morellet et al., 2013; Rivrud et al., 2010; Valderrama-Zafra et al., 2024; van Beest et al., 2011).

The increasing pressure of human activities into natural habitats is altering resource availability, causing habitat loss and disrupting connectivity, with profound impacts on animal movements. Linear (transportation) infrastructure (e.g., roads, fences), forestry, agriculture, hunting, recreation and other human pressures have permeated into natural areas to such an extent that approximately 75% of the Earth’s land surface shows measurable anthropogenic effects (Venter et al., 2016). Human pressures have been summarized into an overall index of human footprint (HFI), including infrastructure, human density, land use change, urbanization, and light pollution (Barnosky et al., 2012; Newbold et al., 2015; Venter et al., 2016). Linear infrastructure can act as barriers, disrupting the connectivity of habitats and, together with changes in resource availability through forestry, agriculture or artificial feeding, alter animal movements and home range sizes (Bischof et al., 2017; Fahrig, 2007; Jerina, 2012; Main et al., 2020; Passoni et al., 2021; Selva et al., 2017). While there is a global trend of animal movements being reduced with increasing human footprint (Main et al., 2020; Tucker et al., 2018), how animals respond to human impact depends largely on the nature of the disturbance and the species life history (Doherty et al., 2021). For example, animals may move further distances in response to direct human disturbances, such as hunting or recreation due to “fear effects” (Doherty et al., 2021).

The scale and ubiquity of disrupted animal movements, in combination with habitat fragmentation, is a major conservation issue worldwide, and in some areas of the world, such as in North America and parts of Asia and Africa, wildlife mainly persists in large, protected wilderness areas away from human habitation that accommodate animals’ spatial needs (Chapron et al., 2014; Packer et al., 2013; Veldhuis et al., 2019). This is especially true for species that have traditionally been the subject of socio-political conflicts when sharing space with humans, like large carnivores, which generally occur at low densities and have large spatial requirements (Bautista et al., 2017). Yet, in the highly fragmented and human- dominated European landscape, large carnivores are currently increasing in numbers and recolonizing their former ranges due to conservation policies, the depopulation of rural areas and increases in forest cover (Chapron et al., 2014; Cimatti et al., 2021; Passoni et al., 2024; Reinhardt et al., 2019). This remarkable comeback was only possible by several behavioral adaptations of large carnivores to live in human-dominated landscapes, e.g., to avoid humans spatially and temporally (de Gabriel Hernando et al., 2020; Lamb et al., 2020; Ordiz et al., 2011). Still, comparative studies evaluating large carnivore behavior along gradients of human disturbance are still lacking.

In this study, we aim to evaluate intraspecific variation in space use patterns of one of the most abundant large carnivores in Europe, the brown bear (*Ursus arctos*), across contrasting environmental and anthropogenic conditions (**Fig 1**). Earlier multi-population studies of brown bear space use from North America have demonstrated substantial intraspecific variation in home range size and overlap, primarily linked to natural food availability (Mcloughlin et al., 2000), but no comparative study has yet tried to determine the drivers of variation in space use patterns across the highly anthropogenic landscape in Europe. With approximately 18,000 individuals, European brown bears are distributed over ten populations that have been increasing or stable during recent years (Kaczensky, 2021). Their range spans from the highly anthropized landscapes in southern and central Europe to the remote boreal forests in northern Finland. Similar to other large carnivores, brown bears act as mobile links in natural ecosystems, playing a significant role in ecosystem dynamics by facilitating seed dispersal and plant regeneration (García-Rodríguez et al., 2021a; García- Rodríguez et al., 2021b; Hämäläinen et al., 2017; Steyaert et al., 2019), protecting plants from herbivorous insects (Grinath et al., 2015), shaping ungulate prey densities (Swenson et al., 2007; Tallian et al., 2021), providing a nitrogen influx into riparian forests (Deacy et al., 2017; Helfield and Naiman, 2006) or removing carrion (Krofel et al., 2012). However, some studies suggest that human-induced altered space use patterns disrupt the role brown bears play in European ecosystems (Diserens et al., 2020; Kuijper et al., 2016).

**Fig. 1.**
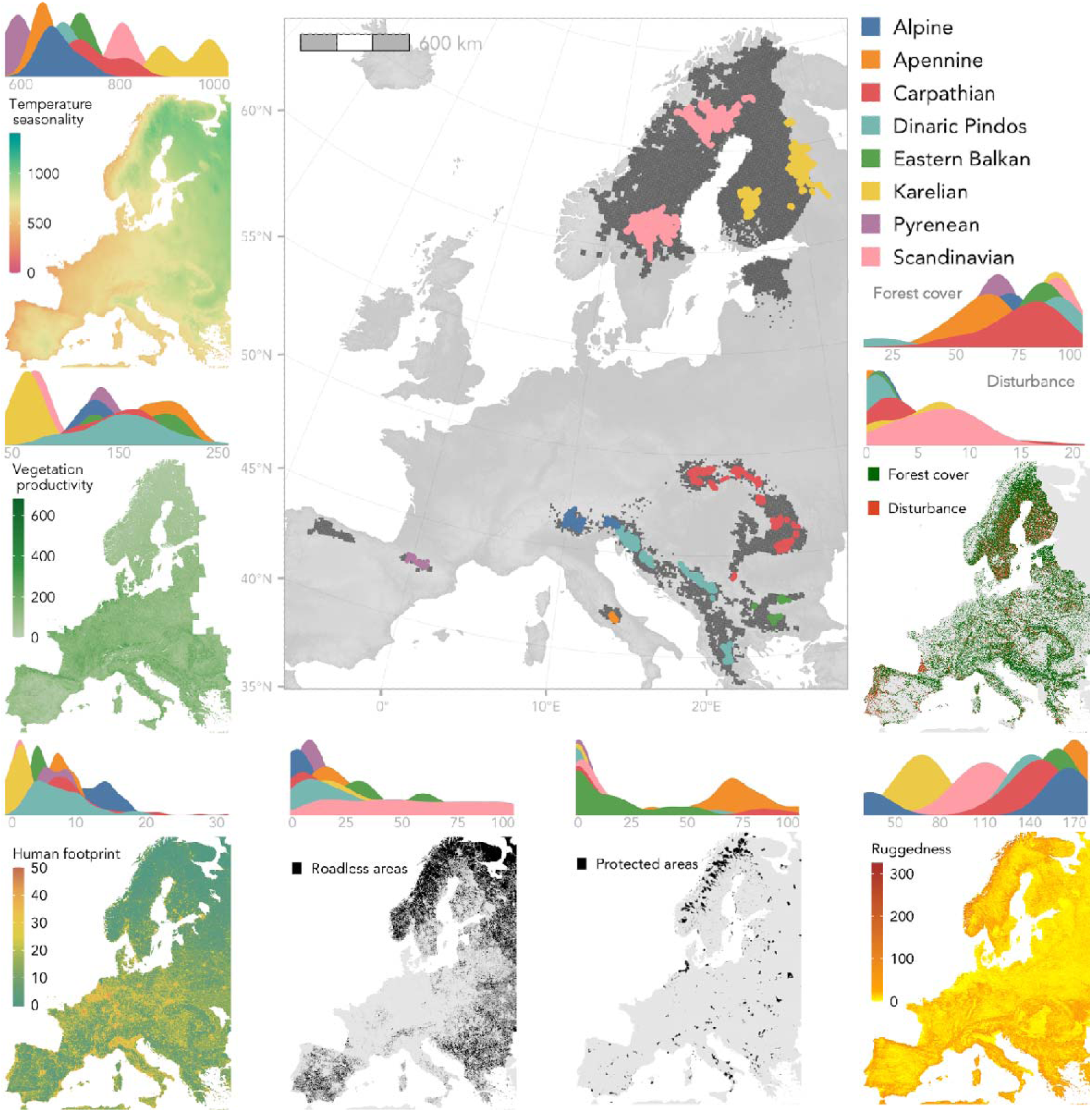
Within the BEARCONNECT initiative, we compiled GPS movement data from eight of the ten extant brown bear populations in Europe. Using data from the summer growing season (May - September) we estimated individual home ranges (main panel, composite home ranges colored by population) which covered a substantial amount of the current permanent occurrence range of the brown bear in Europe (main panel dark gray, Kaczensky 2021). For each home range we extracted the median temperature seasonality (Bio4 in WorldClim, Fick & Hijmans 2017), median annual vegetation productivity (Copernicus 2020), proportion of forest cover and disturbance (Senf & Seidl 2021), median human footprint index (Venter et al. 2016), proportion of roadless (Ibisch et al. 2016) and strictly protected areas (World database on Protected Areas), and median terrain ruggedness index (calculated from a European digital elevation model). Density plots show the distribution of covariates for each population.

We used 752 annual brown bear movement trajectories from the BEARCONNECT movement database (bearconnect.org) to analyze the space use of brown bears across the species’ European geographical range. We delineated annual space use at three distinct spatio-temporal scales during the growing season (May - September), when bears are active and not hibernating: home range size, 10-day long-distance displacement distances, and routine 1-day displacement distances. We summarized a suite of spatially explicit covariates at the home range scale (**Fig. 1**) to evaluate hypotheses concerning the drivers of intraspecific variation in space use.

First, we expected that bears could satisfy their energetic requirements over less space in resource-rich and stable environments (Mcloughlin et al., 2000; Morellet et al., 2013). We therefore predicted smaller home range sizes and shorter movement distances in areas of higher annual vegetation productivity and lower temperature seasonality. In addition, early successional or seral forests after an anthropogenic or natural forest disturbance may provide high quantity, pulsed food resource such as berries due to their open canopy and increased light availability at the ground level (Larsen et al., 2019; Lodberg-Holm et al., 2019; Nielsen et al., 2004). We, therefore, predicted that bears would occupy smaller ranges when they encompass a higher proportion of recently disturbed forests in early successional stages (until 9 years after disturbance (Larsen et al., 2019).

Second, we expected that anthropogenic pressures, as summarized by the human footprint index, would significantly affect bear space use through barrier effects and altered food availability (i.e., increase in food availability and predictability through agriculture or artificial feeding) (Main et al., 2020; Mumme et al., 2023; Tucker et al., 2018). We predicted that bears would occupy smaller home ranges and move less in areas with a higher human footprint index. Conversely, we expected bears to move more in areas of higher forest cover, indicating contiguous, natural habitats with fewer barriers (Cimatti et al., 2021).

Third, we expected that area-based conservation measures, i.e., protected areas or areas with restricted access, have the potential to maintain the ecological integrity of habitats, especially in anthropized landscapes and, thus, may sustain animal movements (Brennan et al., 2022; Hofmann et al., 2021; Jones et al., 2018). We tested the effect of two area-based conservation measures, the proportion of protected areas (WDPA Consortium, 2004) as well as the proportion of roadless areas (Ibisch et al., 2016), within a bear’s home range on bear space use. While roadless areas represent lands relatively undisturbed by humans and are clearly associated with fewer barriers and fragmentation, and increased landscape permeability (Bischof et al., 2017; Lamb et al., 2018), the effects of protected areas are less clear (Geldmann et al., 2013), especially in Europe where the size of protected areas may be either too small to contain brown bear home ranges (Woodroffe and Ginsberg, 1998), where protected areas may be situated in resource poor habitats (Joppa and Pfaff, 2009), or where protected areas are hotspots for recreational activities (Schägner et al., 2016), all of which could ultimately promote animal movements. We predicted that bears move more when their home ranges encompass larger proportions of roadless areas, while the proportion of protected areas could promote or restrict movements.

Last, we accounted for topography, country-level bear population density (Kaczensky, 2021), and for the sex of the individual. We expected that bears inhabiting home ranges with on average more undulated and rugged terrain would move slower and occupy smaller ranges as compared to bears inhabiting flatter terrain. We also expected that the size of home ranges would decrease with increasing population density (Dahle and Swenson, 2003). Given the mating system and social organization of brown bears, we expected males to roam over larger areas than females as shown by previous studies (Nilsen et al., 2005b; Steyaert et al., 2012). We further accounted for variation in bear space use patterns among populations across Europe, which could not be attributed to the main covariates but constitute unexplained variation in space use between populations.

In addition to these main covariates, we also explored the effect of age, reproductive class and artificial feeding on space use, using subsets of our data for which such metadata were available, and we tested for sex-specific responses to our most influential environmental covariates.

## 2. Methods

### 2.1 Compilation and filtering of movement data

As part of a large collaborative Biodiversa project (BEARCONNECT), we compiled a database of all available GPS location data sets from brown bears across Europe. In total, movement data was compiled from eight of the ten extant European brown bear populations (Kaczensky, 2021), spanning 13 countries between 2002 and 2018 representing 615 unique individuals monitored over 1411 tracking years. First, we split our data per individual and year (hereafter referred to as annual bear track). Because brown bears cease movement during winter hibernation, our analysis focused on the summer growing season (i.e., from May to September), assuming that all individuals were completely active during these months. The length of GPS tracking, the GPS sampling interval, and the success rate of GPS locations varied greatly. We therefore only included annual bear tracks that covered at least 100 out of 153 days of the active season (i.e., 3.5 months out of 5 months) and we resampled trajectories to a 1-day resolution, i.e., retaining 1 location/24hrs in an attempt to obtain unbiased and comparable data using the R package *amt* (Signer et al., 2018). Our final dataset included space use information from 752 annual bear tracks from 370 individuals (211 females and 159 males) monitored for 1-13 years, though sample sizes varied substantially among populations (**Fig. 2**).

**Fig 2.**
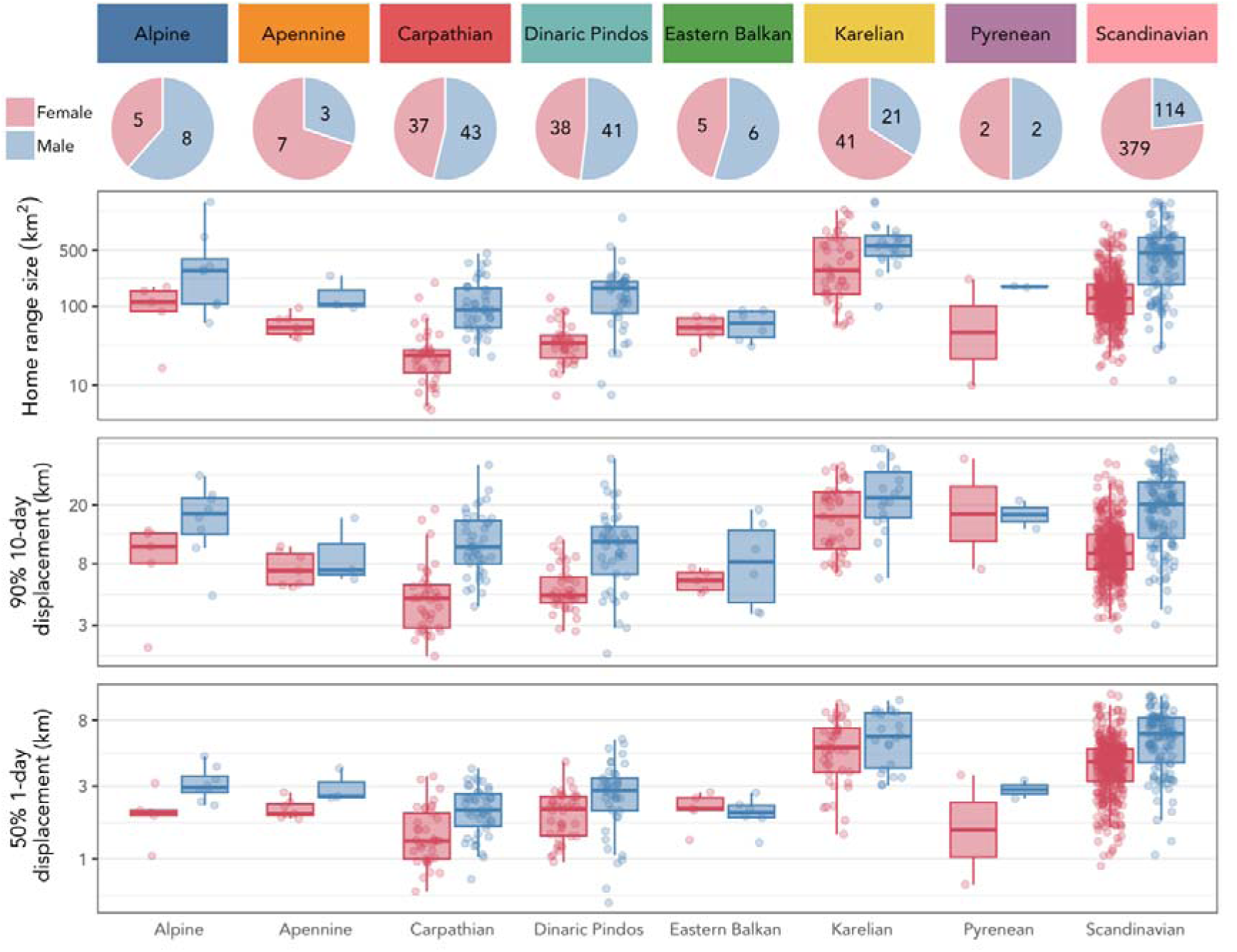
Sex-specific sample sizes (n annual bear tracks) and distribution of space use metrics: home range sizes (km^2^), 10-day and 1-day displacement distances (km), collected for eight European brown bear populations. Male bears generally moved more and occupied larger home ranges than females. All space use metrics were log-transformed for analyses but Y – axis labels are back-transformed to the km^2^/km scale for interpretation.

### 2.2 Space use metrics

We calculated and summarized bear ranging behavior at three distinctive spatio-temporal scales during the summer growing season, leading to one value for each metric per annual bear track: home range size (representing overall space requirements), long distance 10-day displacement (representing long-distance movements), and average 1-day displacement distance (representing routine daily movements).

#### 2.2.1 Home range size

We estimated year-specific home range size for each annual bear track (i.e., the spatially constrained area used by an individual during the summer growing season period between May and September). To do so, we used the time-local convex hull estimator at the 95% isopleth (T-LoCoH, R library ‘tlocoh’ (Lyons et al., 2013), which progressively aggregates local convex hulls to build the home range polygon. We incorporated time and used the adaptive LoCoH method in our home range estimates. Due to the variability of movement patterns among the monitored animals, we selected the time parameter ‘s’ and the parameter for defining neighboring points ‘a’ using the graphical tools available in the T-LoCoH software (see the user guidelines by (Lyons et al., 2013) for more details on the parameters’ selection). Given that topography can affect home range estimates (Morellet et al., 2013), we calculated home range sizes taking into account the three-dimensionality of land topography. For this, we used the Elevation map of Europe (1km grid) from the European Environmental Agency, based on the global digital elevation model derived from GTOPO30 (downloaded from https://www.eea.europa.eu/), and the ‘surfaceArea’ function from the R package ‘sp’ to calculate home range size (**Appendix S1)**. To avoid including annual bear tracks with non-sedentary spatial behavior (i.e., bears that did not occupy a home range during the growing season) in our analyses, we removed from our dataset 12 annual bear tracks that showed directed long-distance dispersal or atypical ranging behavior (**Appendix S2)**.

#### 2.2.2 10-day & 1-day displacement distances

We calculated displacement distances at 10- and 1-day temporal scales, as Euclidean distances between consecutive GPS locations of each annual bear track using the R package *adehabitatLT* (Calenge, 2006). At the 10-day scale, we calculated the 90^th^ percentile of displacement distances for each annual bear track, representing long-distance movements over long temporal scales. At the 1-day scale, we calculated the 50^th^ percentile of displacement distances for each annual bear track, representing routine daily movements (Tucker et al., 2018).

### 2.3 Environmental covariates

We obtained spatial layers of eight environmental covariates related to food availability, human impact, area-based conservation measures, and topography (**Fig. 1**). We extracted and summarized the values of all pixels falling within each bear home range. Thus, every annual bear track was characterized by a single value for each of the tested covariates. We also obtained other covariates that could influence bear space use, in particular, a bear population density index (*2.3.7* and *2.3.8*).

#### 2.3.1. Temperature seasonality

To account for latitudinal differences in seasonality we extracted the median temperature seasonality within each bear’s home range from the WorldClim version 2.1 climate data (Fick and Hijmans, 2017). Temperature seasonality (BIO4 layer) is a measure of temperature change over the course of the year and is calculated as the standard deviation of the annual range in temperature, where larger values represent more seasonal environments and lower values more stable environments with continuous food availability. The BIO4 layer represents the average temperature seasonality for the years 1970-2000. We used the BIO4 layer at a spatial resolution of 2.5 minutes (∼21km) as we wanted to capture large scale latitudinal and altitudinal differences.

#### 2.3.2. Vegetation productivity

To capture differences in food resource availability, we extracted the length of the growing season (in days) and the total vegetation productivity from the European Environmental Agency’s Vegetation Phenology and Productivity (HR-VPP) product suite (Copernicus, 2020; Tian et al., 2021), that represents the growing season integral as the sum of all daily Plant Phenology Index values (PPI, range 0-3) between the dates of the season start and end in a given year. Higher values indicate higher annual productivity and/or longer growing seasons. These maps had a spatial resolution of 100 m and were compiled from finer resolution Sentinel-2A and Sentinel-2B satellite products (10 m resolution and 5-day revisit time). Unfortunately, maps were only available starting in 2017, matching with the last year of our bear movement database and we used the 2017 map as representative for our 16-year dataset of monitoring (2002 - 2018). While we cannot detect site-specific effects of interannual variation in vegetation productivity on movement, our approach does capture large scale latitudinal variation in vegetation productivity. Because total vegetation productivity and length of the growing season are inherently correlated, we selected total vegetation productivity as a better representation measure for resource availability forward into our modeling approach.

#### 2.3.3. Human footprint index

The human footprint index (HFI) map was downloaded from the global map of anthropogenic impact at a 1 km resolution that combines multiple sources of anthropogenic disturbance, including human population density, built-up areas, nighttime lights, crop- and pasture land use, roads, railroads, waterways (Venter et al., 2018; Venter et al., 2016). HFI ranges from 0 to 50, with increasing values indicating high levels of human impact. We extracted HFI values and calculated the median HFI within each home range.

#### 2.3.4. Total forest cover and forest disturbances

Senf and Seidl (2021) published a map of Europe’s forests and identified forest disturbances from USGS Landsat satellite data across Europe between 1986 – 2016 at a spatial resolution of 30 m. The spatial product identifies forest cover (binary variable) and if and when a forest was disturbed between 1986 - 2016. Forest disturbances were defined as cleared forest patches due to either anthropogenic (e.g., forest management and logging) or natural causes (e.g., windfall, fire, bark beetle outbreak, (Senf and Seidl, 2021). Within a bear’s annual home range, we calculated the proportion of forest cover and the proportion of forests in early successional stages after a disturbance, i.e., disturbances occurring within 9 years before the year a bear track was recorded (Larsen et al., 2019). We excluded the year of disturbance for two reasons: 1) structural disturbance of the site likely does not lead to increased food availability in the first year, 2) we wanted to mitigate any effects of forestry activity on animal movement.

#### 2.3.5. Roadless and protected areas

We downloaded the protected areas with the highest degree of IUCN protection (i.e., Strict Nature Reserve – Ia, Wilderness Area – Ib, and National Park – II) as shape files from the World Database on Protected Areas (WDPA) and World Database on Other Effective Area-based Conservation Measures (WD-OECM, https://www.protectedplanet.net). These IUCN categories represent areas of high ecological integrity but potentially with high human disturbance through recreation (Jones et al., 2018). We calculated the proportion of the home range that was covered by protected areas. We further used the global map of roadless areas (shape file), representing areas relatively free of barriers (Ibisch *et al*. 2016, downloaded from http://www.roadless.online/). Roadless areas were defined as land units that were at least 1 km away from any kind of mapped roads (Ibisch et al., 2016). We calculated the proportion of the home range that was covered by roadless areas.

#### 2.3.6. Terrain ruggedness

We calculated the terrain ruggedness index TRI (Riley et al., 1999), a measure of topographic heterogeneity, using the Elevation map of Europe from the European Environmental Agency (https://www.eea.europa.eu/) at a 1 km resolution. TRI ranges from level terrain (values of 0 - 80), nearly level (81 - 116), slightly rugged (117 - 161), intermediately rugged (162 - 239), moderately rugged (240 - 497), to highly rugged terrain (>498). We calculated the median TRI of all values falling within a home range.

#### 2.3.7. Population density index

We assigned to each annual bear track a country-specific bear population density index (individuals/km^2^), which we estimated using permanent occurrence maps (Kaczensky, 2021), at a 10×10 km ETRS89-LAEA Europe grid scale (effective 2012 – 2016), and the country-specific population size published by the IUCN Red List of Threatened Species 2016. Specifically, we divided the population size by the population-specific area of permanent occurrence in each country. Our country-specific bear population density index ranged from 0.007 bears/km^2^ in the Apennine to 0.075 bears/km^2^ in Romania and Croatia (**Table S1**). Substantial within-country spatial variation in bear population density remains which we could not account for because to date, there are no bear population density maps available.

#### 2.3.8 Additional covariates explored in the Appendix

*Artificial feeding*: For each country, we extracted information about the use of artificial feeding (binary: yes/no) as a management tool from Bautista *et al*. (2017). Artificial feeding can constrain animal movement (Selva et al., 2017), however, no comprehensive data are available for Europe on when, where, and how much food is supplied to wildlife. Therefore, we contrasted bear space use in countries where artificial feeding is or is not allowed (**Table S1, Appendix S3**). *Age class*: Age class is known to affect bear space use as young dispersing bears, in particular males, often roam over larger areas (Dahle and Swenson, 2003). While we excluded bear tracks showing directed dispersal, young bears can also show home range shifts over time. Bear age was not recorded for all bears in a standardized way. Therefore, we categorized a subset of bears for which we had some age information (**Table S7**) as either subadult (1 – 4 years of age) or adult (> 4 years of age), and tested whether subadult males or females would occupy larger home ranges or move over longer distances (**Appendix S4**).

### 2.4. Statistical analyses

We fitted full Bayesian linear mixed effects models for each of the log-transformed space use metrics: home range size, the 90th percentile of 10-day displacement distances, and the 50th percentile of 1-day displacement distances. We accounted for the main effects of annual vegetation productivity, temperature seasonality, HFI, the proportion of forest cover, the proportion of early successional forest, the proportion of protected and roadless areas, terrain ruggedness, sex, and population density. We further incorporated random intercepts for population and individual identity. We tested sex-specific responses to the most relevant environmental gradients (see **Appendix S5)**. Our models did not suffer from multicollinearity, as indicated by a variance inflation factor < 2 for all comparisons. All models were fitted with a Gaussian family with the R package *brms* (Bürkner 2017), running four chains over 4000 iterations with a warmup of 2000 and a thinning interval of 2. The model inference was based on 4000 posterior samples and had satisfactory convergence diagnostics with ^R < 1.01 and effective sample sizes > 1000. Posterior predictive checks recreated the underlying Gaussian distribution well and did not show signs of heteroscedasticity. We report the median as a measure of centrality and 89% credible intervals, calculated as equal tail intervals, as a measure of uncertainty (Kruschke, 2014; McElreath, 2020). Data and code to reproduce the analysis are available via the Open Science Framework (Hertel, 2024).

## 3. Results

Intraspecific variation of brown bear space use in Europe was evident on all spatio-temporal scales. Home range sizes varied from 72 to 260 km^2^ (1^st^ and 3^rd^ quartile, median = 129 km^2^), 10-day long-distance displacements from 7 to 16 km (median = 10 km), and daily displacements from 2.5 to 5.4 km (median = 3.9km; **Fig 2**). The smallest estimates were recorded in the Carpathian and Dinaric Pindos populations, and the largest in the Karelian population (**Fig 2**). Males occupied larger home ranges and moved more in all populations, except for the Eastern Balkan (**Fig 2, Table S2**). The three space use metrics were positively correlated (**Fig S1**). Across populations, European brown bears occupied a wide range of environments (**Fig 1).** For example, bears in the Karelian and Scandinavian populations experienced the lowest vegetation productivity, highest temperature seasonality, occupied home ranges with the highest proportions of early successional forest cover and experienced the lowest HFI (**Fig 1**, **Table S3**).

### 3.1. Drivers of space use patterns

Intraspecific variation in brown bear space use patterns was explained by sex, proxies of resource availability and human pressure. Vegetation productivity, the proportion of early successional forests, HFI, and sex had a significant effect on brown bear space use across spatio-temporal scales, while the proportion of protected area had a weak effect on long-distance displacements only (**Fig 3**). At the shortest temporal scale, i.e., during routine daily displacements, the proportion of forest cover and terrain ruggedness additionally affected bear movements. All models explained a good amount of variance in the data but generally fixed covariates performed better at explaining intraspecific variation in movement over longer time scales, i.e., on the home range (marginal R^2^ = 0.44) and 10-day scale (marginal R^2^ = 0.36), than on the daily short time scale (marginal R^2^ = 0.25).

**Fig 3.**
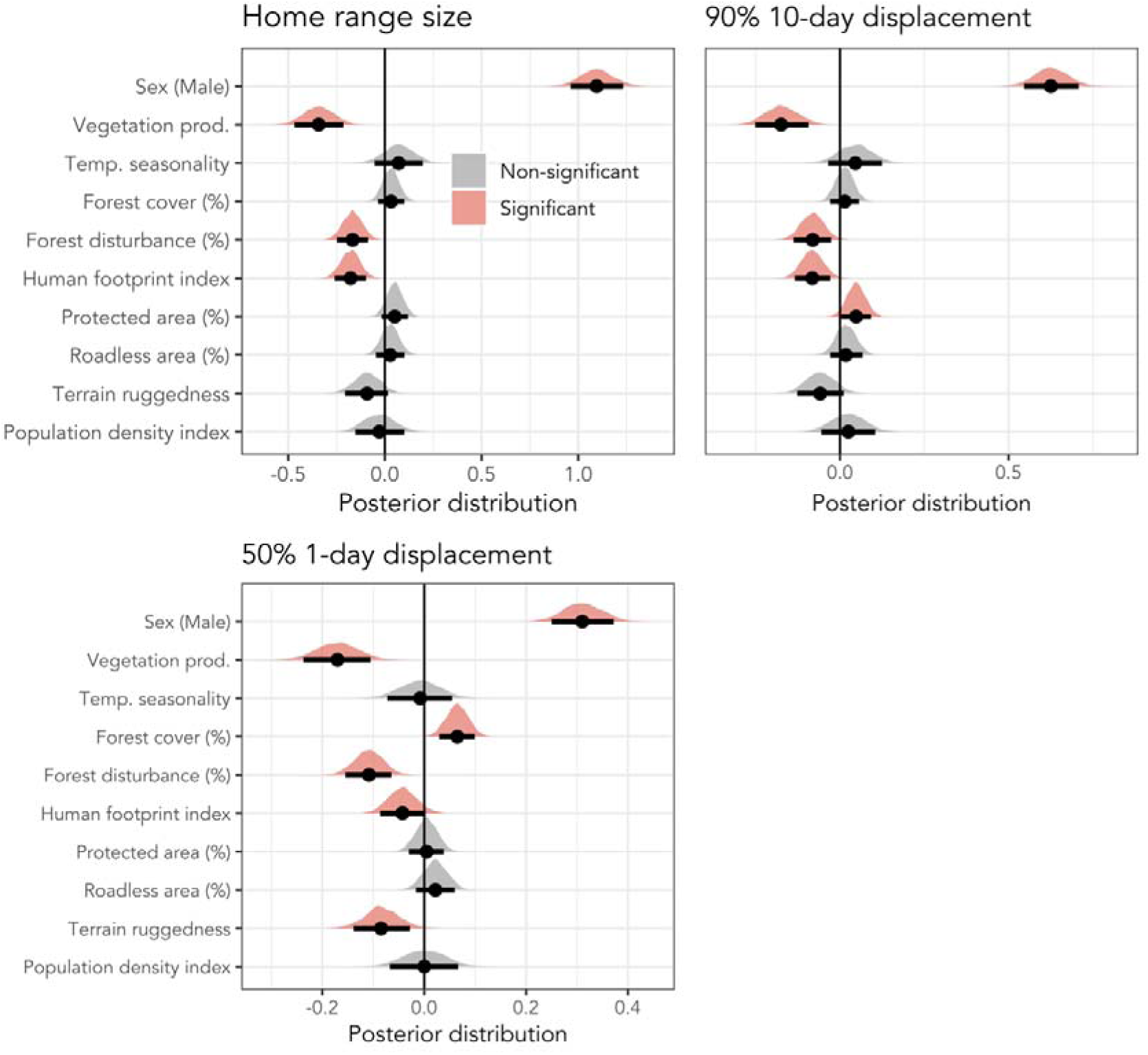
Coefficients plots showing effect sizes of covariates on space use metrics of brown bears across Europe. We found consistent effects of sex, human footprint index, and the proportion of forest disturbances within a bear’s home range on bear movement and space use. See also **Table S5** for all model coefficients.

In line with our first prediction, two proxies of resource availability - vegetation productivity and the proportion of disturbed forest in early successional stages - were negatively correlated with bear space use at all temporal scales, while temperature seasonality had no significant effect **(Fig 3).** Specifically, in areas of low vegetation productivity, home ranges were five times larger than in areas of high productivity (**Fig 4a**) and bears moved twice as far both at the 10-day and 1-day scale (predicted 10-day displacement at low productivity (45) = 12 km vs high productivity (250) = 5 km; predicted 1-day displacement 3.6 km vs 1.5 km, **Fig 4d & g**). In addition, the proportion of forest disturbances in a home range was also related to bear space use at all spatio-temporal scales. Bears occupied home ranges with a median proportion of 5% [range = 0% - 21%] of forests that were disturbed within last 9 years. Bears occupied smaller home ranges and moved over shorter distances when their home range encompassed a greater proportion of recent forest disturbances (**Fig 4b, e & h**, predicted home range size at 0% = 128 km^2^ vs at 15% = 63 km^2^; predicted 10-day displacement at 0 % = 11 km vs at 15% = 7.5 km; predicted 1-day displacement at 0 % = 3.4 km vs at 15% = 2 km). General forest cover only affected 1-day displacements, with bears moving longer distances in areas with more forest cover (predicted 1-day displacement in areas with 13% forest cover = 2 km vs 100% forest cover = 3.1km). (**Fig. 3**).

**Fig 4.**
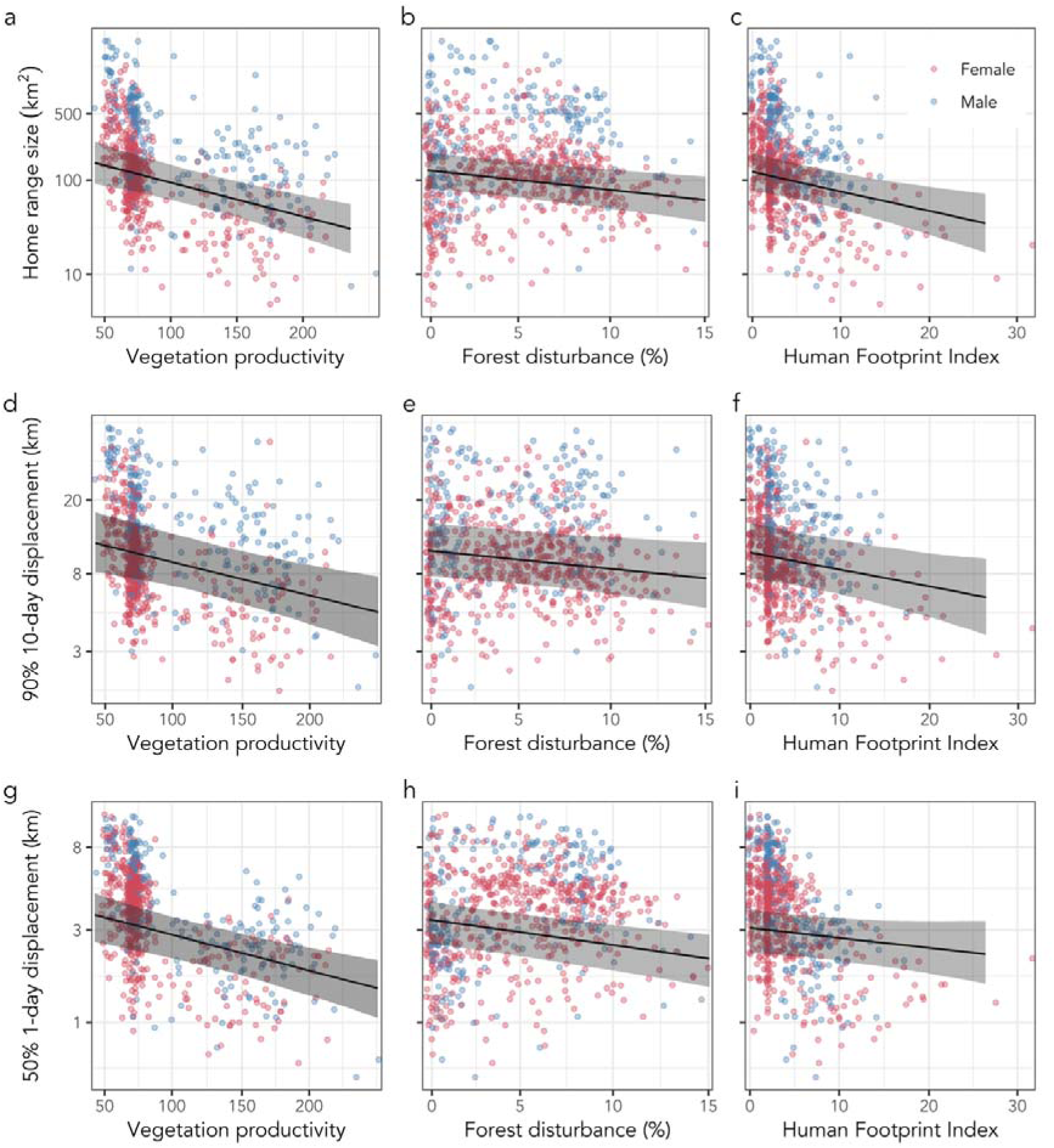
Effect sizes of vegetation productivity, the proportion of early successional forest, and human footprint index on home range size, long-distance displacements, and routine displacements of brown bears across Europe. These three covariates significantly affected space use across spatio-temporal scales. Space use decreased with increasing vegetation productivity (a, d, g), with an increasing proportion of recent forest disturbances (i.e., early successional forest) in a bear’s home range (b, e, h), and with increasing median human footprint index in the home range (c, f, i).

Human footprint index strongly shaped intraspecific variation in brown bear space use (**Fig 3**), supporting our second hypothesis. Bears occupied home ranges with a median HFI between 0 and 32 (median = 2.5, **Table S3**), i.e., from natural to highly modified environments, wherein human footprint consistently shaped bear space use across all scales. In general, with increasing HFI, bears formed smaller home ranges and moved less (**Fig 3**). This reduction was most apparent at larger spatio-temporal scales. For example, bears in areas with high HFI occupied home ranges a quarter the size of those in low HFI areas (predicted home range size at HFI 32 = 28 km^2^ vs at HFI 0 = 123 km^2^, **Fig 4c**). Similarly, 10- day long-displacements in areas with low HFI were twice as long compared to highly disturbed areas (predicted 10-day displacement at HFI 32 = 5 km vs at HFI 0 = 10 km, **Fig 4f**). At shorter time scales the effect was less strong but still apparent – predicted daily displacements in undisturbed landscapes (HFI 0) were 3.1km, while in disturbed landscapes (HFI 32) routine movements were reduced to 2.1km (**Fig 4i**).

Most home ranges overlapped only marginally with protected areas (median = 2.2% [range = 0%, 100%]) and we found weak and mixed support that area-based conservation measures affect brown bear space use (**Fig 3**). Long-distance displacements increased with the proportion of protected area in a bears’ home range (predicted 10-day displacement at 0% protected area = 9.2km vs. at 100% protected area = 12km) but home range size and routine daily movements were unaffected by the proportion of protected area. Although bear home ranges readily overlapped with roadless areas (median = 37% [range = 0%, 100%), intraspecific variation in bear space use patterns was not influenced by the proportion of roadless areas in a home range.

Finally, terrain ruggedness affected bear space use at short temporal scales only, with shorter daily displacements in more rugged terrain. Country-level bear population density index, did not affect home ranges or movement distances (**Fig 2**). Sex-specific differences in bear space use were strong and evident across spatio-temporal scales (**Fig 3**). Across populations, males formed home ranges that were three times the size of females (304 vs 101 km^2^) and male long-term displacements covered twice the distance (18 km for males compared to 9.5 km for females). However, routine daily displacements were similar for both sexes (4 and 3 km, resp.). Despite the profound sex differences in space use, we found no evidence for sex-specific responses to environmental covariates (**Appendix S5**): male and female home range size decreased in a similar fashion with increasing vegetation productivity, proportion of early successional forest, and human footprint.

### 3.2. Population-level differences in space use

For all three space use metrics, substantial variation remained that could not be explained by fixed covariates. Between-population differences, i.e., bears belonging to a given population behaving in a similar fashion and different from bears in other populations, accounted for 22%, 30%, and 25% of the variance in home range size, 10-day long-distance displacements, and 1-day routine displacements, respectively. Additionally, between-individual differences accounted for 51%, 35%, and 10% of variance. The total explained variance including fixed and random effects (conditional R^2^) was 81% for home range size, 68% for long-time displacement, and 54% for daily displacements (**Table S5**). Based on the posterior distribution of the random intercept for study population (after accounting for fixed covariates, **Fig 5**), we found that bears from the Carpathian, Dinaric Pindos, and Eastern Balkan populations showed limited space use across all spatio-temporal scales: they occupied the smallest home ranges (75km^2^) and moved over shortest distances (long distance = 7.5km, routine daily displacements = 2.5km). Bears from the Italian Alpine population occupied large home ranges (165km^2^) and showed long-distance displacements (12km) but moved over intermediate distances on a routine daily basis (2.9km). Bears from the Apennine and Karelian populations occupied large home ranges (125 km^2^) and moved over long distances across scales (long distance = 10km, routine daily displacements = 3.5km). Last, Scandinavian bears occupied intermediate home range sizes (93km^2^) and moved over intermediately long distances (8.5km), but showed the longest movements on a daily scale (3.6km). Estimates for the Pyrenean population should be treated with caution because of the limited available movement data, stemming from only three individuals included in our final dataset that were reintroduced and tracked post-release (**Table 1**).

**Fig. 5.**
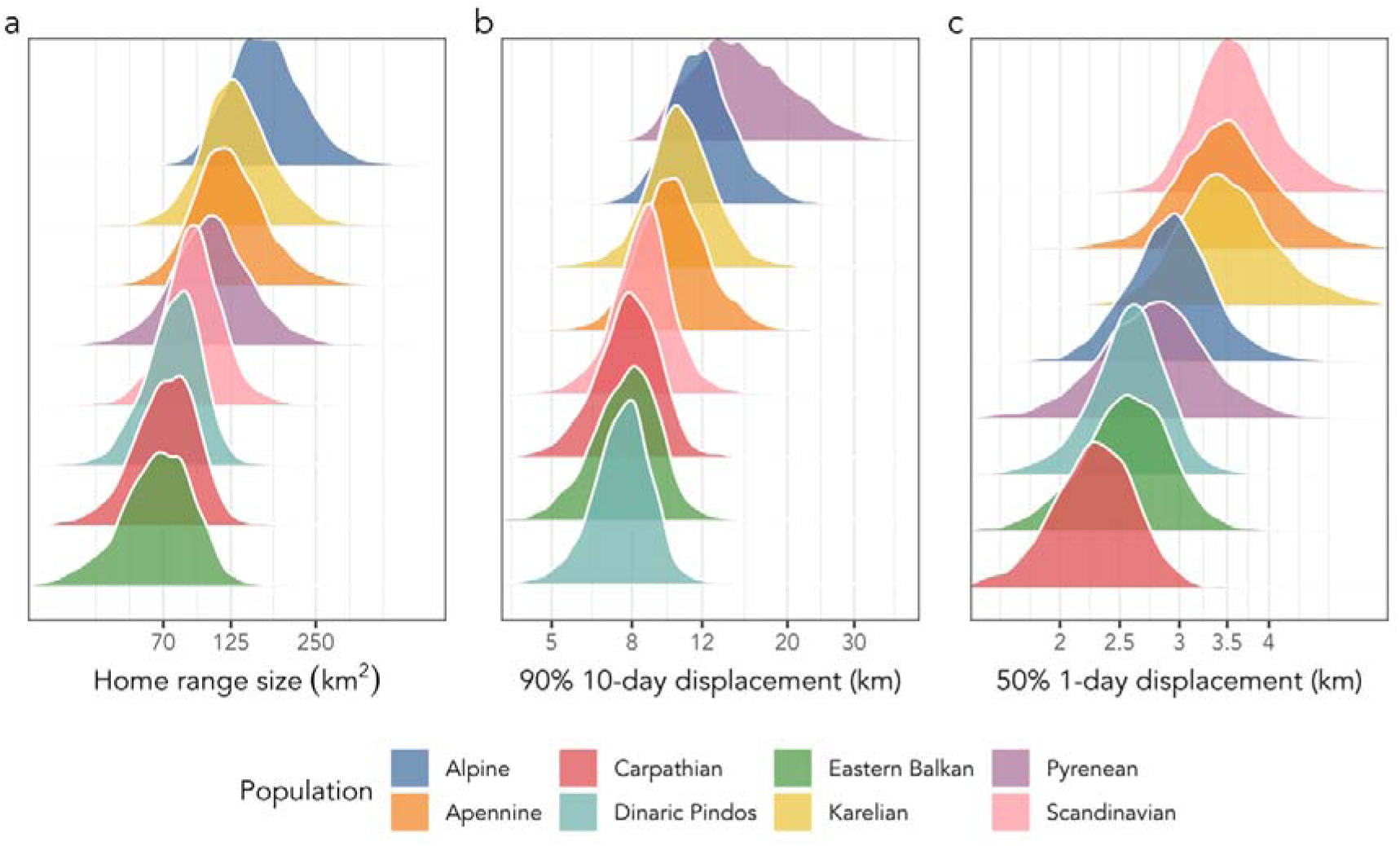
Between-population differences after accounting for fixed covariates are shown as the posterior distribution of the random intercept for each study population. Between population differences in (a) home range size ranged from 70 – 165 km^2^, in (b) 10-day displacements ranged from 7.5 – 15km, and (c) in 1-day displacements ranged from 2.3 – 3.6 km.

### 3.3. Artificial feeding

Nine out of 14 countries included in our study generally provided artificial food to bears (**Table S1**), representing 211 of 751 annual bear tracks from the Carpathian, Dinaric Pindos, Eastern Balkan and Karelian populations. We had no information on the amount, spatial, or temporal distribution of artificial feeding at the individual home range level. While the raw data suggested smaller home ranges and shorter daily movements for bears that were artificially fed as compared to ones that were not fed, after controlling for all other covariates, model coefficients suggested a positive effect (**Appendix 3, Table S6**).

## 4. Discussion

We found that brown bear space use patterns across the European continent were jointly governed by resource availability, human impact, and terrain but were not affected by area-based conservation measures. Specifically, bears occupied smaller home ranges and moved less in areas of higher vegetation productivity or where recently disturbed, early successional forests provide abundant food, supporting our first hypothesis. Increasing human footprint restricted bear space use while increasing proportions of forest cover promoted movement, supporting our second hypothesis and corroborating earlier findings of reduced mammalian movements in areas with high human pressures at global scales (Doherty et al., 2021; Main et al., 2020; Mumme et al., 2023; Tucker et al., 2018). Our findings suggest that human footprint hinders landscape permeability for brown bears on a continental scale. Contradicting to these findings and to our third hypothesis, the amount of roadless area in bear home ranges did not affect space use, potentially because of the high fragmentation and small size of roadless areas in Europe (Ibisch et al., 2016). Alternatively, bears might show restrict space use in areas of high human footprint because these areas provide abundant food through artificial feeding, croplands, orchards, beehives or trash (Bautista et al., 2021). Human impacts, including human footprint, forest disturbances, artificial feeding, and recreation in protected areas, are ubiquitous across Europe and we here provided the first comprehensive overview on how anthropogenic effects govern brown bear spatial behavior.

### 4.1. Resource availability shapes space use: the role of vegetation productivity, forestry, and artificial feeding

Brown bear populations in south-eastern Europe occupied the smallest home ranges and moved over shorter distances, while populations in Fennoscandia occupied the largest ranges. This marked intraspecific variability seems to be the outcome of different cost-to-benefit ratios of moving and acquiring resources at varying levels of human disturbance and resource availability along the species’ distribution range. In line with previous studies, a gradient of natural variation in plant food availability drove intraspecific variation in space use (Mcloughlin et al., 2000). Home ranges were smaller and movement distances shorter in areas of higher vegetation productivity, a commonly used proxy of forage availability. Vegetation productivity thereby not only accounts for herbaceous foods but also affects the carrying capacity of ungulate prey. Food resources are more limiting at the northern latitudes due to the gradual decline in the length of the growing season and greater temperature seasonality, and consequently, animals need to move over larger areas to find sufficient food resources (Mcloughlin et al., 2000; Morellet et al., 2013). Therefore, bears living at the northern edge of the distribution range appear to compensate for the lower vegetation productivity by foraging over larger areas. However, unlike for vegetation productivity, we did not find a direct link between temperature seasonality and bear space use.

Additionally, we found that the proportion of recently disturbed forest patches led to a reduction in movement and smaller home ranges. Such early successional forests often offer clustered and abundant food resources, such as ants or berries, and bears may be able to satisfy caloric needs over less area and with shorter movement distances. While we could not quantify whether these disturbances were anthropogenic (i.e., forestry) or natural (i.e., bark beetle, wind falls) disturbances (Senf and Seidl, 2021), it has been estimated that 95% of all forest disturbances in Europe are due to forestry (Curtis et al., 2018). It is noteworthy that our inference on how recent forest disturbances affect bear space use comes primarily from the Scandinavian, Karelian (boreal needleleaf forest), and the Carpathian population (broadleaf forests), as the proportions of recent forest disturbances were small in all other populations (**Fig 1, Table S3**). This aligns with Sweden and Finland being the biggest timber producing industries in Europe, where estimated 2% of forest area is harvested every year (Ceccherini et al., 2020). While several studies have evaluated the relationship between forest clearings, food abundance, and bear space use within populations (Frank et al., 2015; Larsen et al., 2019; Nielsen et al., 2004), we here provide the first generalizable evidence that forest disturbance may affect species movements across larger spatial scales and biomes. Future research should evaluate whether more generalizable patterns regarding, e.g., successional forest patch size or distribution, on animal space use emerge.

We expected that space use would additionally be linked to the exploitation of anthropogenic food resources in human-dominated landscapes, such as agricultural fields or artificial feeding sites, as animals need to travel less to find food (Doherty et al., 2021). Although a reduction in home ranges and movement distances associated with artificial feeding has been previously shown in brown bears and other mammal species (Jerina, 2012; Selva et al., 2017), we did not confirm this reduction at the continental scale. However, our inference was limited by the fact that we only had country-level binary information on whether artificial feeding was used or not as a management tool, which is too coarse to demonstrate causal links between artificial feeding and space use across populations (see **Appendix S3**). We still assume that artificial feeding drives differences in space use, as bears in the Carpathian, Dinaric Pindos and Easter Balkan populations occupied the smallest home ranges and moved least, while these populations are also the ones where artificial feeding is a prevalent management tool (Bautista et al., 2017; Selva et al., 2017). In summary, the link between bear space use and food availability was strong and supported through multiple pathways, namely, vegetation productivity, forest disturbances, and potentially artificial feeding. Climate change is predicted to alter vegetation and fruit-based food availability across the latitudinal gradient of Europe. For example, in Spain warming temperatures have been linked to shifts in brown bear diet away from boreal and temperate food items (Penteriani et al., 2019). And in Fennoscandia, winter warming and freezing events have been suggested to reduce berry crops, in particular on early successional forest stands that are lacking a protective canopy cover (Hertel et al., 2018). Elsewhere, phenological synchronization of food items has been observed, disrupting the seasonal succession of food availability (Deacy et al., 2017). Our results suggest that climate induced shifts in food availability and abundance may result in changes of brown bear space use patterns.

### 4.2. Human footprint restricts space use

Brown bears in Europe occupied home ranges with a median human footprint ranging from 0 to 32, which aligns with the human footprint range occupied by other large wildlife, such as red deer (Main et al., 2020; Mumme et al., 2023; Tucker et al., 2018). Bears in central and southern Europe were exposed to higher levels of human pressures than bears in Fennoscandia (**Fig 1**), with bears in the Italian Alps occupying home ranges with the highest human footprint (Passoni et al., 2024). Across Europe, bears moved less and occupied smaller home ranges in areas with a higher human footprint. This reduction suggests that some of the human pressures included in the human footprint index, e.g., human settlements or high-traffic roads (Selva et al., 2011), can act as barriers to bear movements at large spatio-temporal scales. Our study is the first to provide generalizable evidence from multiple populations that brown bear space use is affected by human pressures across biomes and environmental conditions. Landscapes with higher human footprint may also provide clustered, high-caloric food, e.g., in agricultural fields or garbage dumps, again modifying space use in human-dominated landscapes via resource availability (Doherty et al., 2021). In line with these findings, a higher proportion of forest cover in a bear’s home range led to longer routine daily movements, suggesting that more contiguous forest cover with fewer edges promote bear movements (Cimatti et al., 2021).

### 4.3. Inconclusive effect of area-based conservation measures

Brown bear space in Europe was largely unaffected by area-based conservation measures, i.e., the proportion of protected and roadless areas within a bears’ home. The only detectable effect was that long-distance displacements were longer in more protected areas. However, the median proportion of protected areas within a home range was only 2%, and bears in the Apennines were the only ones overlapping primarily with a protected area. Given the lack of overlap of bear home ranges with protected areas, we could not conclusively test the effect of protected areas on bear movement behavior. We suggest that protected areas in Europe are too small to encompass the spatial needs of brown bears and to impact their movement behavior.

Finally, brown bear space use was not affected by population density index, even though previous studies have documented its effects on mammal ranging behavior (Mcloughlin et al., 2000). However, the population density data utilized in our study were measured at a coarse country level scale and may not accurately reflect local densities. Higher local population densities are likely supported by higher resource availability and our results suggest that space use is restricted by food availability and not population density per se.

### 4.4. Potential consequences of altered space use for connectivity and ecosystem services

While the return of large carnivores in Europe can be hailed a conservation success, it also highlights that large carnivore species must have strong behavioral adaptability in order to coexist with humans (Gaynor et al., 2019; Gaynor et al., 2018; Lamb et al., 2020). These behavioral adjustments can come at a survival cost in human-dominated landscapes (Cosgrove et al., 2018; Lamb et al., 2016; Oriol-Cotterill et al., 2015). In addition, our observed disruption in long-distance 10-day displacements in particular, may have implications for population expansion and demographic connectivity, potentially impeding dispersal or mate searching behavior (Bartoń et al., 2019), and thereby promoting genetic isolation (Bischof et al., 2017; Epps et al., 2005). Especially for small and isolated populations (e.g., in Southern Europe), decreasing genetic diversity, inbreeding and inbreeding depression have been found, which can compromise population viability (Benazzo et al., 2017; De Barba et al., 2010; Palazón et al., 2012). For the small bear populations in the Apennine and the Pyrenees, which have long been isolated and have no prospect for connectivity with other populations, actions tailored at supporting movements and range expansion are critical to promote population growth for their recovery (Kervellec et al., 2023; Maiorano et al., 2019). Where populations are in close proximity but barriers such as high traffic roads and settlements restrict movement, such as between the Dinaric Pindos and Carpathian populations in Serbia, or the Alpine and Dinaric Pindos populations in the Alps, corridors mitigating human impacts may aid in establishing inter-population connectivity (Bogdanović et al., 2023; Peters et al., 2015). The special protection of long-distance dispersers in particular, and of wide-ranging movement in general has also been suggested as a conservation measure to support population connectivity, particularly needed in highly-modified landscapes like Europe (Bartoń et al., 2019). Ultimately recovering bear populations harbor the potential for increasing human-wildlife conflict (Bautista et al., 2017). This is particularly true in areas where bears have been formerly extirpated and people have forgotten how to coexist with them and where bears share space with humans (Passoni et al., 2024; Tosi et al., 2015).

Bears play an important role in the functioning of terrestrial and aquatic ecosystems and as connectivity umbrella species (Diserens et al., 2020; Dutta et al., 2023; Helfield and Naiman, 2006) and the human-induced reduction of their movements could have cascading impacts on many ecosystem processes and services (Cosgrove et al., 2018; Doherty et al., 2021). As mobile omnivores (frugivores to a great extent), that travel between habitats and ecosystems with varying levels of human footprint, they may be instrumental in shaping trophic interactions and rewiring food webs (Bartley et al., 2019; Grinath et al., 2015; Ripple et al., 2014). Bears appear to be effective as connectivity umbrellas for several other coexisting mammals in Fennoscandia, as well as in smaller populations in Eastern Europe and Dinaric Pindos (Dutta et al., 2023), emphasizing the value of the species in anthropized regions. Although the functional role of bears in European ecosystems with higher human footprint can be questioned, studies have shown that, when the appropriate management measures are taken, their role as seed dispersers can be preserved, even in areas with high human pressure (García-Rodríguez et al., 2021b). However, management practices such as artificial feeding are likely to disrupt seed dispersal processes and predation effects on ungulate populations (Kuijper et al., 2016). To our knowledge, the only region in Europe where any measurable top-down regulatory effects by brown bears on ungulates have been demonstrated is Scandinavia (Støen et al., 2022; Tallian et al., 2021), where both vegetation productivity and human footprint are generally low, no supplementary food is provided, and space use and mobility are higher than in central or southern Europe. Such evidence suggests potentially profound ecosystem-wide consequences from reduced space use, in Central and Southern Europe’s human-dominated landscapes. Further research on how space use affects the role bears play in their ecosystem is needed. The implementation of effective measures to preserve animal movements in areas with a high human footprint will be key for the connectivity of recovering brown bear populations in Europe. Our research emphasizes the role of food availability and forest disturbances, in further restricting animal movement, and demonstrates the value of forest cover to promote animal movements.

## Conclusion

The observed intraspecific variability in brown bear space use was governed by the different conditions in resource availability and human footprint across the brown bear distribution range in Europe. Bears reduced space use in areas of increased vegetation productivity, in areas with more forest disturbances, and higher human footprint. These results support the mounting evidence that point to a global restructuring of animal movement caused by the intensification of human activities (Doherty et al., 2021; Main et al., 2020; Mumme et al., 2023; Tucker et al., 2018). With the current expansion of large carnivores in the highly fragmented European landscape, reducing the negative impacts of humans on animal movement will be of key importance to ensure the successful future conservation of these populations and the functioning and resilience of the ecosystems they inhabit.

## Acknowledgements

BearConnect is an initiative for large scale data integration and mobilization for brown bears in Europe for conservation. It is currently self-funded by BearConnect project partners and supported by close collaboration with researchers and stakeholders. BearConnect was initially funded by BiodivERsA (Horizon 2020 ERA-NET COFUND) through the 2015-2016 BiodivERsA COFUND call, with national funders ANR (ANR16-EBI3-0003), NCN (2016/22/Z/NZ8/00121), BMBF DLR-PT (01LC1614A), CCCDI – UEFISCDI (BiodivERsA3-2015-147-BearConnect (96/2016)), and RCN (269863). AGH has received funding from the German Science Foundation (HE 8857/1-1). AP was funded by the National Science Centre in Poland (project PRELUDIUM 2022/45/N/NZ8/04127).

The research was supported by the following entities

<u>Italy:</u> GPS locations in the Apennine bear population were collected in a research project funded by a private U.S. donor through the Wildlife Conservation Society, LM and PC were funded by the European Union - NextGenerationEU National Biodiversity Future Center. Access to the GPS telemetry data for the brown bears expanding from Trentino, Italy was provided by the Edmund Mach Foundation.

Poland: the project GLOBE (POL-NOR/198352/85/2013) funded by the Norway Grants under the Polish-Norwegian Research Programme operated by the National Center for Research and Development, and Tatra National Park brown bear conservation plan.

Romania: Romanian Ministry of Research, Innovation and Digitalization funds - PN23090304 (12N/01.01.2023);

Scandinavia: The Scandinavian Brown Bear Research Project is funded by the Swedish Environmental Protection Agency and the Norwegian Environmental Protection Agency; Serbia: The research was supported by the Serbian Ministry of Education, Science and Technological Development (451–03–68/2022–14/200178).

<u>Slovakia:</u> GPS telemetry study of bears in Slovakia was supported by Dr. Joachim and Hanna Schmidt Stiftung für Umwelt und Verkehr, Germany, and the project ITMS 26220120069 under the Operational Programme Research and Development funded by the European Regional Development Fund.

<u>Slovenia:</u> Brown bear telemetry studies in Slovenia were financed by the European Union, Ministry of the Environment and Spatial Planning of Slovenia, Slovenia Research and Innovation Agency and Slovenian Environment Agency (projects J4-7362, LIFE13 NAT/SI/000550, P4-0059)

## Data availability statement

Data and code to reproduce the analysis have been archived in the OpenScience Framework under the accession number https://doi.org/10.17605/OSF.IO/XH39F. Restrictions apply to the raw GPS data; these can be obtained from the corresponding authors upon reasonable request.

