## Supplementary material for "Human footprint and forest disturbance reduce space use of brown bears (*Ursus arctos*) across Europe"

**Online supplementary material**

- **Table S1. Overview of population specific metadata: Population size, hunting and artificial feeding regime**
- **Table S2. Population specific median and range of response variables.**
- **Table S3. Population specific covariate values**
- **Table S4. Testing for nonlinear effects**
- **Table S5. Model coefficients of selected models presented in the main text.**
- **Figure S1**. **Correlation of response variables**
- **Figure S2**. **Correlation of predictor variables**
- **Appendix 1. Incorporation of the three-dimensional surface into our home range estimates**
- **Appendix 2. Identification of non-sedentary individuals**
- **Appendix 3. Effects of supplementary feeding on movement**
- **Appendix 4. Age effects on movement**
- **Appendix 5. Sex-specific effects of predictors**

**Table S1.** Overview of population and country specific population size, and information on harvest and artificial feeding (for bears as target species).

| **Population** | **Country** | **Population size (IUCN 2018)*** | **Area of occurrence (km^2^, 2012 - 2016)**** | **Population density** | **Bear harvest** | **Supplementary feeding** |
| --- | --- | --- | --- | --- | --- | --- |
| Alpine | Italy | 58 | 5308 | 0.011 | no | no |
| Apennine | Italy | 58 | 8000 | 0.007 | no | no |
| Carpathian | Poland | 100 | 4316 | 0.023 | no | yes |
| Carpathian | Romania | 6075 | 81564 | 0.074 | yes | yes |
| Carpathian | Slovakia | 1261 | 20597 | 0.061 | yes | yes |
| Dinaric-Pindos | Bosnia-Herzegovina | 1000 | 17943 | 0.056 | yes | yes |
| Dinaric-Pindos | Croatia | 937 | 12355 | 0.076 | yes | yes |
| Dinaric-Pindos | Greece | 475 | 15977 | 0.030 | no | no |
| Dinaric-Pindos | Serbia | 120 | 8012 | 0.015 | no | yes |
| Dinaric-Pindos | Slovenia | 564 | 9660 | 0.058 | yes | yes |
| Eastern Balkan | Bulgaria | 421 | 17709 | 0.024 | yes | yes |
| Karelian | Finland | 1780 | 128629 | 0.014 | yes | yes |
| Pyrenean | France/Spain | 43 | 3600 | 0.012 | no | no |
| Scandinavian | Sweden | 2782 | 222679 | 0.012 | yes | no |

*https://www.iucnredlist.org/species/41688/144339998#assessment-information

**Kaczensky, Petra et al. (2021). Distribution of large carnivores in Europe 2012 - 2016: Distribution maps for Brown bear, Eurasian lynx, Grey wolf, and Wolverine [Dataset]. Dryad. https://doi.org/10.5061/dryad.pc866t1p3

**Table S2.** Population and sex-specific sample sizes and median space use metrics (for females / males, resp.).

| **Population** | **N_individuals_ (female/male)** | **N_bear-year tracks_ (female/male)** | **Median home range size (km^2^)** | **Median 90% 10-day displacement (km)** | **Median 50% 1-day displacement (km)** |
| --- | --- | --- | --- | --- | --- |
| Alpine | 3 / 7 | 5 / 8 | 113 / 282 | 10 / 18 | 2 / 2.9 |
| Apennine | 6 / 3 | 7 / 3 | 54 / 106 | 7 / 7 | 2 / 2.6 |
| Carpathian | 25 / 35 | 37 / 43 | 25 / 91 | 5 / 10 | 1.3 / 2.1 |
| Dinaric Pindos | 31 / 35 | 38 / 41 | 35 / 168 | 5 / 11 | 2.1 / 2.8 |
| Eastern Balkan | 4 / 6 | 5 / 6 | 54 / 63 | 6 / 8 | 2.1 / 2 |
| Karelian | 24 / 12 | 41 / 21 | 284 / 574 | 17 / 23 | 5.3 / 6.2 |
| Pyrenean | 2 / 1 | 2 / 2 | 115 / 177 | 25 / 18 | 2.1 / 2.8 |
| Scandinavian | 116 / 60 | 379 / 114 | 125 / 466 | 9 / 21 | 4.3 / 6.5 |

**Table S3.** Population specific covariate values (median [minimum, maximum]).

| **Population** | **Seasonality** | **Vegetation productivity** | **Forest cover (%)** | **Forest disturbance (%)** | **HFI** | **Roadless areas (%)** | **Protected areas (%)** | **Terrain ruggedness index** |
| --- | --- | --- | --- | --- | --- | --- | --- | --- |
| Alpine | 667 [628, 734] | 134 [102, 194] | 68 [39, 79] | 1 [1, 2] | 12 [7, 17] | 3 [0, 26] | 0 [0, 21] | 162 [29, 179] |
| Apennine | 642 [637, 655] | 192 [163, 216] | 63 [41, 77] | 0 [0, 1] | 8 [5, 10] | 16 [5, 38] | 70 [34, 100] | 170 [163, 174] |
| Carpathian | 728 [669, 833] | 155 [92, 216] | 80 [25, 98] | 3 [0, 21] | 9 [3, 32] | 10 [0, 96] | 2 [0, 98] | 145 [114, 167] |
| Dinaric Pindos | 686 [645, 747] | 157 [81, 252] | 88 [13, 100] | 1 [0, 5] | 7 [2, 20] | 15 [0, 95] | 0 [0, 100] | 141 [120, 170] |
| Eastern Balkan | 715 [698, 753] | 174 [122, 205] | 84 [73, 95] | 1 [0, 2] | 5 [4, 8] | 34 [13, 76] | 3 [0, 54] | 159 [139, 167] |
| Karelian | 965 [867, 1011] | 65 [44, 85] | 89 [81, 97] | 6 [0, 9] | 2 [0, 7] | 23 [0, 83] | 0 [0, 42] | 68 [39, 95] |
| Pyrenean | 594 [568, 617] | 136 [128, 173] | 67 [61, 71] | 1 [0, 1] | 7 [5, 10] | 9 [6, 11] | 0 [0, 6] | 172 [163, 175] |
| Scandinavian | 809 [770, 1019] | 72 [49, 93] | 90 [26, 98] | 7 [0, 17] | 2 [0, 8] | 50 [0, 100] | 3 [0, 98] | 108 [41, 140] |

**Table S4.** Nonlinear effects

| **Home Range** | **Fixed** | **waic** | **waic_se** | **marginalR2** | **conditionalR2** |
| --- | --- | --- | --- | --- | --- |
| Full linear effects | hfi_median + sex + vegetationproductivity + prop_protected + prop_roadless + prop_forest + prop_disturbed_forest + population_density | 1243 | 43 | 0.44 [0.33, 0.55] | 0.81 [0.78, 0.83] |
| Polynomial hfi | hfi_median + I(hfi_median,2) | 1246 | 43 | 0.45 [0.34, 0.56] | 0.81 [0.79, 0.83] |
| Polynomial vegetation | vegetationproductivity + I(vegetationproductivity,2) | 1242 | 42 | 0.44 [0.32, 0.55] | 0.81 [0.79, 0.84] |
| Polynomial forest disturbance | prop_disturbed_forest + I(prop_disturbed_forest,2) | 1225 | 42 | 0.49 [0.38, 0.59] | 0.82 [0.79, 0.84] |
| **50% daily displacement** | **Fixed** | **waic** | **waic_se** | **marginalR2** | **conditionalR2** |
| Full linear effects | hfi_median + sex + vegetationproductivity + prop_protected + prop_roadless + prop_forest + prop_disturbed_forest + population_density | 828 | 41 | 0.27 [0.14, 0.41] | 0.55 [0.5, 0.6] |
| Polynomial hfi | hfi_median + I(hfi_median,2) | 824 | 41 | 0.29 [0.15, 0.43] | 0.55 [0.5, 0.6] |
| Polynomial vegetation | vegetationproductivity + I(vegetationproductivity,2) | 830 | 41 | 0.26 [0.13, 0.41] | 0.55 [0.5, 0.6] |
| Polynomial forest disturbance | prop_disturbed_forest + I(prop_disturbed_forest,2) | 824 | 43 | 0.31 [0.14, 0.45] | 0.55 [0.5, 0.6] |
| **90% 10-day displacement** | **Fixed** | **waic** | **waic_se** | **marginalR2** | **conditionalR2** |
| Full linear effects | hfi_median + sex + vegetationproductivity + prop_protected + prop_roadless + prop_forest + prop_disturbed_forest + population_density | 805 | 40 | 0.38 [0.28, 0.47] | 0.68 [0.64, 0.72] |
| Polynomial hfi | hfi_median + I(hfi_median,2) | 807 | 40 | 0.37 [0.28, 0.47] | 0.68 [0.64, 0.72] |
| Polynomial vegetation | vegetationproductivity + I(vegetationproductivity,2) | 800 | 40 | 0.36 [0.28, 0.46] | 0.69 [0.65, 0.73] |
| Polynomial forest disturbance | prop_disturbed_forest + I(prop_disturbed_forest,2) | 805 | 40 | 0.4 [0.3, 0.5] | 0.69 [0.65, 0.73] |

**Table S5.** Median (measure of centrality) and 89% equal tails interval (measure of uncertainty) of model coefficients

|  | **Home range size** | **10-day displacement** | **1-day displacement** |
| --- | --- | --- | --- |
| **Fixed effects** |  |  |  |
| Intercept | 18.44 [18.11, 18.77] | 9.16 [ 8.93, 9.44] | 7.98 [ 7.76, 8.15] |
| Sex (male) | **1.1 [ 0.96, 1.23]** | **0.63 [ 0.55, 0.71]** | **0.31 [0.25, 0.37]** |
| Median HFI | **-0.18 [-0.26, -0.1]** | **-0.08 [-0.13, -0.03]** | **-0.04 [-0.09, 0]** |
| Vegetation prod. | **-0.34 [-0.47, -0.21]** | **-0.17 [-0.25, -0.09]** | **-0.17 [-0.24, -0.11]** |
| Seasonality | 0.07 [-0.05, 0.2] | 0.05 [-0.04, 0.12] | -0.01 [-0.07, 0.05] |
| Protected areas (%) | 0.05 [-0.02, 0.12] | **0.05 [0, 0.09]** | 0 [-0.03, 0.04] |
| Roadless areas (%) | 0.03 [-0.05, 0.1] | 0.02 [-0.03, 0.07] | 0.02 [-0.02, 0.06] |
| Forest cover (%) | 0.03 [-0.04, 0.1] | 0.01 [-0.03, 0.06] | **0.06 [0.03, 0.1]** |
| Disturbed forest (%) | **-0.17 [-0.25, -0.09]** | **-0.08 [-0.14, -0.03]** | **-0.11 [-0.17, -0.06]** |
| Ruggedness | -0.09 [-0.21, 0.02] | -0.06 [-0.13, 0.01] | **-0.08 [-0.14, -0.03]** |
| Population density | -0.03 [ 0.15, 0.1] | 0.02 [-0.06, 0.1] | 0 [-0.07, 0.07] |
| **Random effects** |  |  |  |
| sd_intercept.BearID_ | 0.65 [0.58, 0.72] | 0.36 [0.3, 0.4] | 0.16 [0.09, 0.22] |
| sd_intercept.Population_ | 0.43 [0.33, 1.45] | 0.33 [0.23, 1.12] | 0.24 [0.1, 0.64] |
| residual | 0.47 [0.44, 0.51] | 0.36 [0.34, 0.39] | 0.4 [0.37, 0.43] |
| **Marginal R2** | 0.44 [0.34, 0.54] | 0.36 [0.26, 0.45] | 0.25 [0.12, 0.41] |
| **Conditional R2** | 0.81 [0.79, 0.83] | 0.68 [0.64, 0.72] | 0.54 [0.49, 0.6] |

**Figure S1.** Correlation of response variables

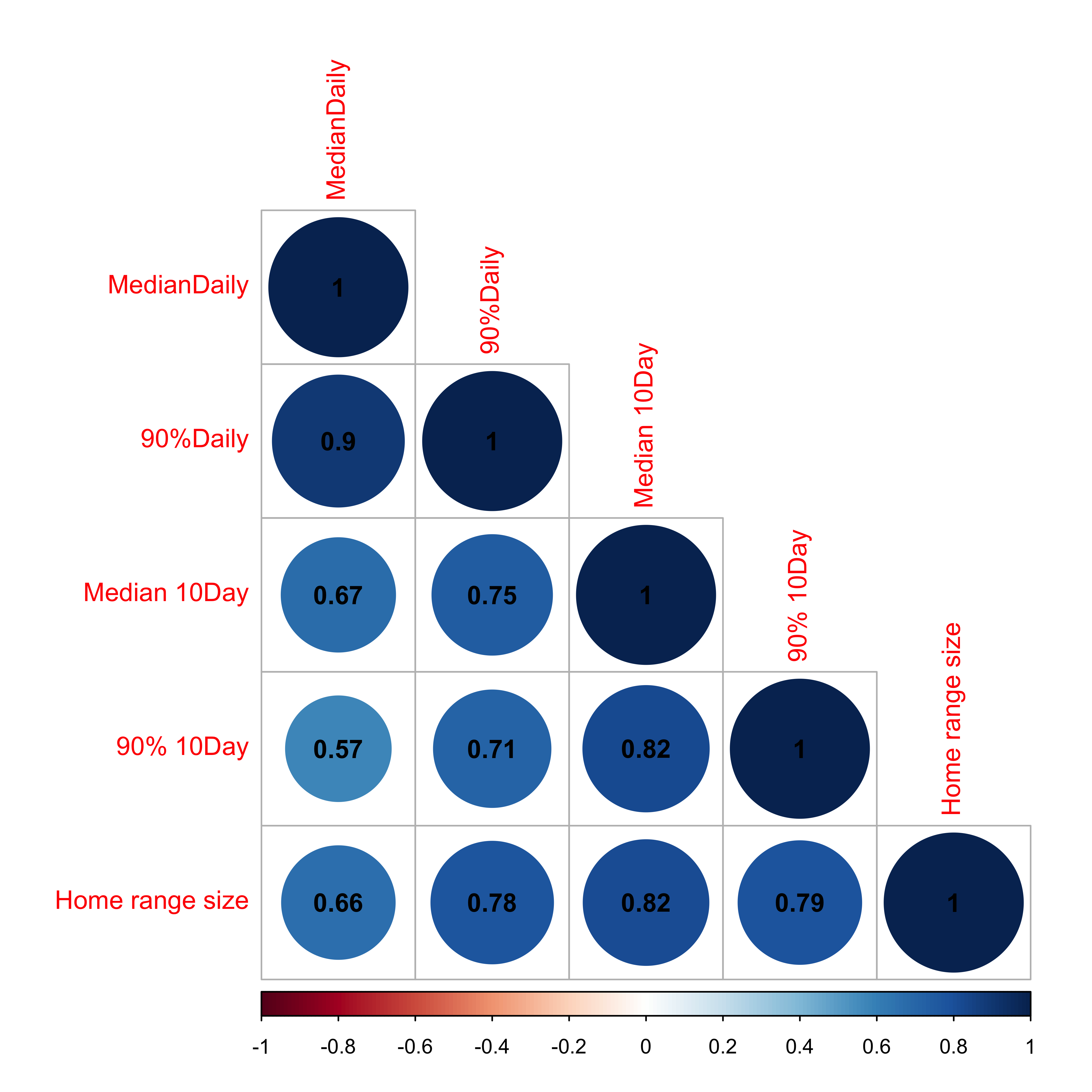

**Figure S2.** Correlation of selected predictor variables. Albeit expectedly substantial correlation of some predictor variables (e.g., seasonality, vegetation productivity, and ruggedness), the variance inflation factor was < 2 for all three models.

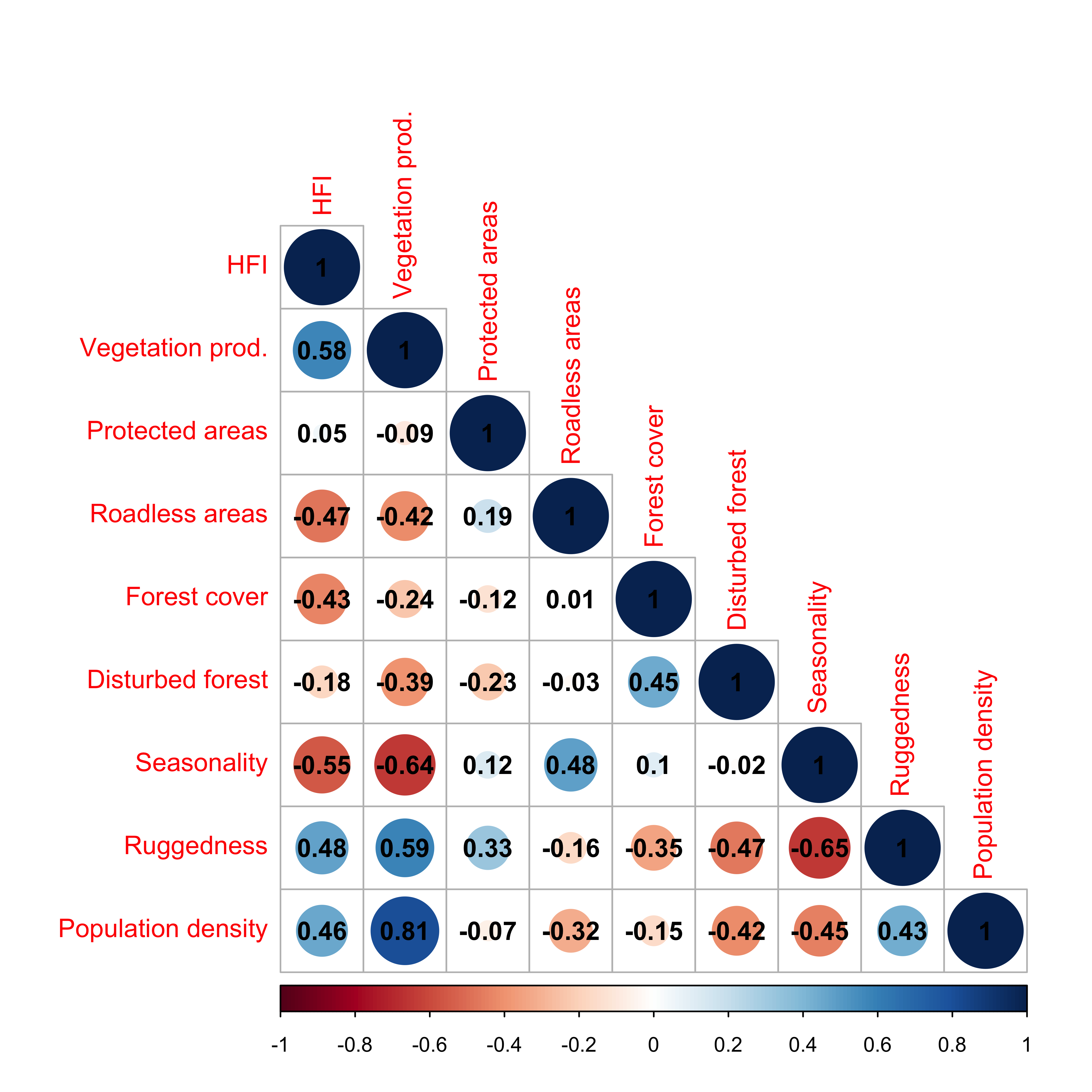

**Appendix 1. Incorporation of the three-dimensional surface into our home range estimates.**

Because home range size can be affected by gradients in elevation or ruggedness of the landscape (Monterroso et al., 2013) and many European brown bear populations inhabit in mountainous areas, we tested whether using home range estimates that account for vertical movement was needed.

To do so, we compared the two and three-dimensional home range estimates using a paired Wilcoxon test and set statistical significance at p = 0.05. Our findings show that home range estimates increased significantly after incorporating the three-dimensional surface (n = 761, V = 0, p < 0.001). Thus, we used the three-dimensional home range estimates in further analyses to ensure the adequate representation of the ranging behavior in our study populations.

**Appendix 2. Identification of non-sedentary individuals**

Our dataset included previously reported cases of non-sedentary individuals (e.g., a long-distance dispersal male from the Carpathian brown bear population, Bartoń et al. 2019). We, therefore, conducted visual inspections of yearly individual tracks during the active period (from May to September) using the net squared displacement (NSD) method to identify directed long-distance displacement behaviors as well as other behavioral pattens typical in non-sedentary individuals. NSD is a convenient metric for identifying specific ranging behaviors, such as home range, dispersal, migration, and nomadic movements, by calculating the distance traveled relative to an origin point (Bunnefeld *et al.* 2011). According to Bunnefeld et al. (2011), animals that restrict their movement to home ranges exhibit an asymptotic NSD curve over time, while dispersers display a sigmoid curve. Our visual inspection identified 12 bear-year tracks with non-sedentary behaviors, including directed long-distance dispersal (e.g. individuals 794_2015, 879_2008, 902_2011 and 5081_2012) and other unclear patterns (Fig. 1). These individuals were excluded from the analyses to ensure that our results exclusively reflected
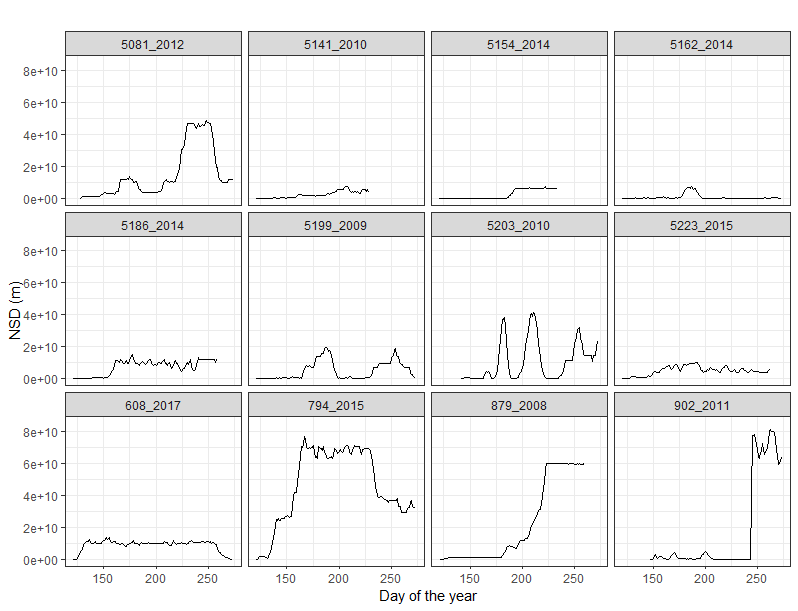
movements within established home ranges.

**Figure S3.** Net squared displacement (NSD) of 12 European brown bears over the active season (from May – day 121 - to September – day 274) who displayed non-sedentary behaviors.

**Appendix 3. The effect of artificial feeding on brown bear space use**

Intentional or unintentional artificial or diversionary feeding of wildlife is a common management strategy. Targeted artificial feeding of bears can have several reasons: a) to keep bears away from human settlements or crops and to reduce conflict (Garshelis *et al.* 2017), b) to attract bears to specific areas for the purpose of wildlife viewing (Penteriani *et al.* 2021), or c) to attract bears during the hunting season (baiting) (Steyaert *et al.* 2014; Kirby, Macfarland & Pauli 2017). In addition, bears often consume food targeted towards other species. Ungulates are often supplementary fed during the winter to buffer natural food shortage and boost survival and reproductive success, elsewhere ungulates are fed to avoid browsing and damage to young trees or agricultural crops (Selva *et al.* 2017; Arnold *et al.* 2018). Bears that use artificial feeding sites have been reported to move less or have smaller home ranges because they can meet their energetic demands by using clustered, high caloric food (Selva *et al.* 2017; Bojarska *et al.* 2019; Penteriani *et al.* 2021) but elsewhere no effects on home range size variability have been found (Todorov 2020). Importantly, even if artificial food is provided, not all bears may use feeding sites.

Across Europe, artificial feeding is common. In our dataset, in nine out of 14 countries provided artificial food targeted for bears, representing 211 of 751 bear monitoring years (**Table S1**). Yet, testing with our dataset whether bears occupy smaller home ranges because of artificial feeding is difficult because information on the amount of food provided is lacking. Looking at the raw data, bears seemed to occupy smaller home ranges (**Fig. S4a**) and in particular moved over shorter daily distances (**Fig. S4c**) where artificial food was provided.

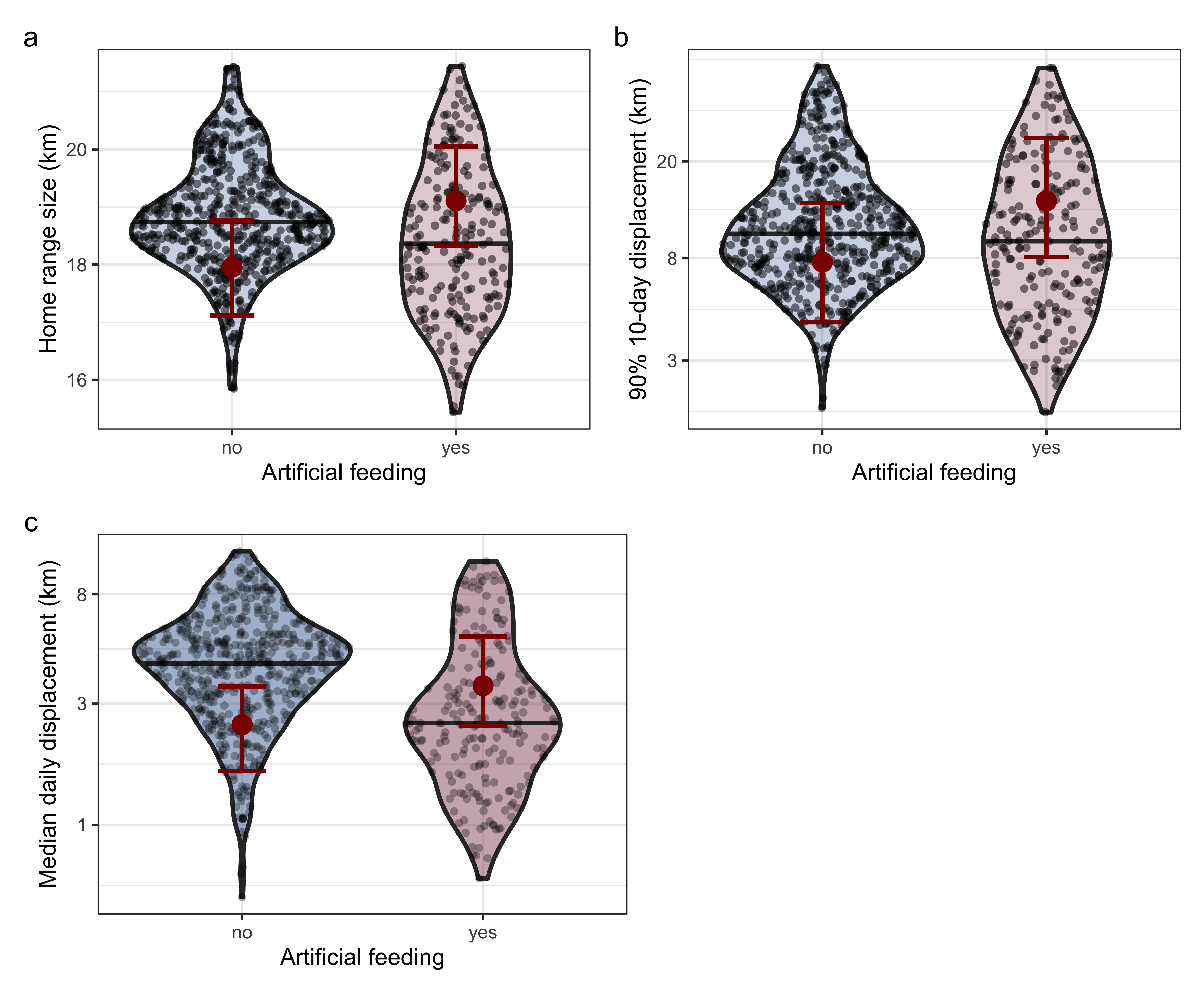

**Figure S4.** Space use of bears in areas with and without bear targeted artificial feeding.

We included artificial feeding as a binary variable (yes/no) into our final models for home range size, 90% 10-day displacement, and median daily displacement. For all three models, the variance inflation factor was > 2 suggesting collinearity with other predictors in the model (VIF for artificial feeding in home range model = 2.17; 10-day displacement = 2.24; daily displacement = 2.06). The effect size was significant in all cases (**Table S6**) in the direction that artificial feeding increases space requirements (estimate and 95% confidence interval shown in red, **Figure S4**). While at first sight, these results are counter intuitive, we have to keep in mind that the models are controlling for the effects of all other variables as well. Therefore, all else being equal, the model predicts larger space use requirements for bears in areas with artificial feeding. We have decided to not include artificial feeding into our final model for two main reasons. First, it has a high variance inflation factor indicating that its colinear with other variables in the model. Indeed, artificial feeding is a common management practice in all south-eastern European countries (with the exception of Greece), in the ranges of the Carpathian, Dinaric-Pindos and Eastern Balkan population. These populations however also have higher vegetation productivity in their home ranges than bears in the other populations which live at higher latitudes or altitudes. Second, the scale at which artificial feeding could be measured (country-wide scale yes/no) is too coarse to know if the bears in our dataset indeed have access to artificial feeding sites and whether they use them. Artificial feeding schemes are also highly variable with regards to how much food is being put out, at what time of year and at how many sites but we were unable to obtain reliable data on the nature of artificial feeding in our dataset. In summary, we think that artificial feeding probably has an effect in space use, either a positive or negative effect but our data did not allow us to draw sound conclusions and we therefore did not include artificial feeding in our main models.

**Table S6.** Model coefficients for home range size, 90% 10-day displacement, and median daily displacement when including artificial feeding regime into the model.

|  | **Home range size** | **10-day displacement** | **Median daily displacement** |
| --- | --- | --- | --- |
| Intercept | 17.8 [16.98, 18.68] | 8.84 [8.3, 9.34] | 7.69 [7.26, 8.09] |
| Sex (male) | 1.12 [0.96, 1.29] | 0.65 [0.55, 0.75] | -0.08 [-0.17, 0] |
| Median HFI | -0.17 [-0.27, -0.07] | -0.08 [-0.14, -0.01] | 0.01 [0, 0.03] |
| Median HFI (2) | - | - | 0.33 [0.25, 0.4] |
| Vegetation productivity | -0.36 [-0.52, -0.21] | -0.18 [-0.28, -0.08] | -0.16 [-0.24, -0.07] |
| Protected areas (%) | -0.02 [-0.11, 0.06] | 0 [-0.05, 0.06] | -0.04 [-0.08, 0.01] |
| Roadless areas (%) | 0.01 [-0.07, 0.1] | 0.02 [-0.04, 0.07] | -0.01 [-0.06, 0.04] |
| Forest cover (%) | -0.06 [-0.15, 0.04] | -0.03 [-0.1, 0.02] | 0.03 [-0.02, 0.08] |
| Disturbed forest (%) | -0.21 [-0.32, -0.11] | -0.1 [-0.17, -0.03] | -0.12 [-0.18, -0.06] |
| Population density | -4.78 [-10.76, 1.11] | -2.11 [-5.68, 1.42] | -0.72 [-3.63, 2.34] |
| Artificial feeding (yes) | **1.2 [0.57, 1.85]** | **0.64 [0.23, 1.05]** | **0.4 [0.09, 0.74]** |

**Appendix 4. Age effects on movement**

Brown bears are generally considered a non-territorial range resident species, where individuals occupy a home range that is regularly traversed in the search for food, mates, and shelter. However, subadult brown bears of both sexes disperse from their natal range (Hansen *et al.* 2021; Thorsen *et al.* 2022) primarily to avoid inbreeding (Støen *et al.* 2006). During dispersal subadult bears show directed long distance movements which leads to larger home range sizes as compared to adults (Dahle & Swenson 2003). We therefore expect age class specific variation in brown bear space use, where subadults occupy larger home ranges and move over longer distances as compared to adults. We additionally expect these effects to be stronger in males than in females due to philopatry in many females (Hansen *et al.* 2021).

Across our dataset, the methodology to estimate age varied from monitoring since birth, over counting the cementum annuli in a bear tooth (Matson *et al.* 1993) to visual age estimation based on mass and structural size. Individual ages were reported as minimum age, age range, estimated age or as a category (adult or subadult). Given the huge discrepancy of age estimates in our dataset we roughly categorized bears as subadult (0 – 4 years) or adult (> 4 years, **Table S7**).

**Table S7.** Population specific number of subadult (age 0-4), adult (age >4) and unknown aged (NA) bear tracks monitored with GPS collars and included in our dataset.

| **Population** | **Female**  **subadult / adult / NA** | **Male**  **subadult / adult / NA** |
| --- | --- | --- |
| Alpine | - / - / 5 | 1 / 3 / 4 |
| Apennine | 1 / 6 / - | 1 / 2 / - |
| Carpathian | 5 / 21 / 11 | 18 / 10 / 15 |
| Dinaric Pindos | 15 / 23 / - | 18 / 23 / - |
| Eastern Balkan | 4 / 1 / - | 6 / - / - |
| Karelian | 12 / 29 / - | 5 / 12 / - |
| Pyrenean | - / 2 / - | - / 2 / - |
| Scandinavian | 101 / 250 / 28 | 46 / 57 / 8 |

We estimated sex-specific age differences in home range size, long distance displacements and routine daily displacements using univariate mixed models accounting for age class and a random intercept for population identity (**Fig S5**). Compared to adult females, subadult females occupied significantly smaller home ranges (estimated adult home range size = 74km^2^, subadult = 54km^2^) and moved over slightly shorter long-distance displacements (adult = 8.5km, subadult = 7.1km), while routine daily displacements were similar for adult and subadult females (both 2.4km). Likewise adult males occupied larger home ranges (adult = 228km^2^, subadult = 117km^2^) and ranged wider, both on the 10-day scale (displacement distance adult = 15km, subadult = 10km) and 1-day scale (displacement distance adult = 3.7km, subadult = 2.5km) than subadult males.

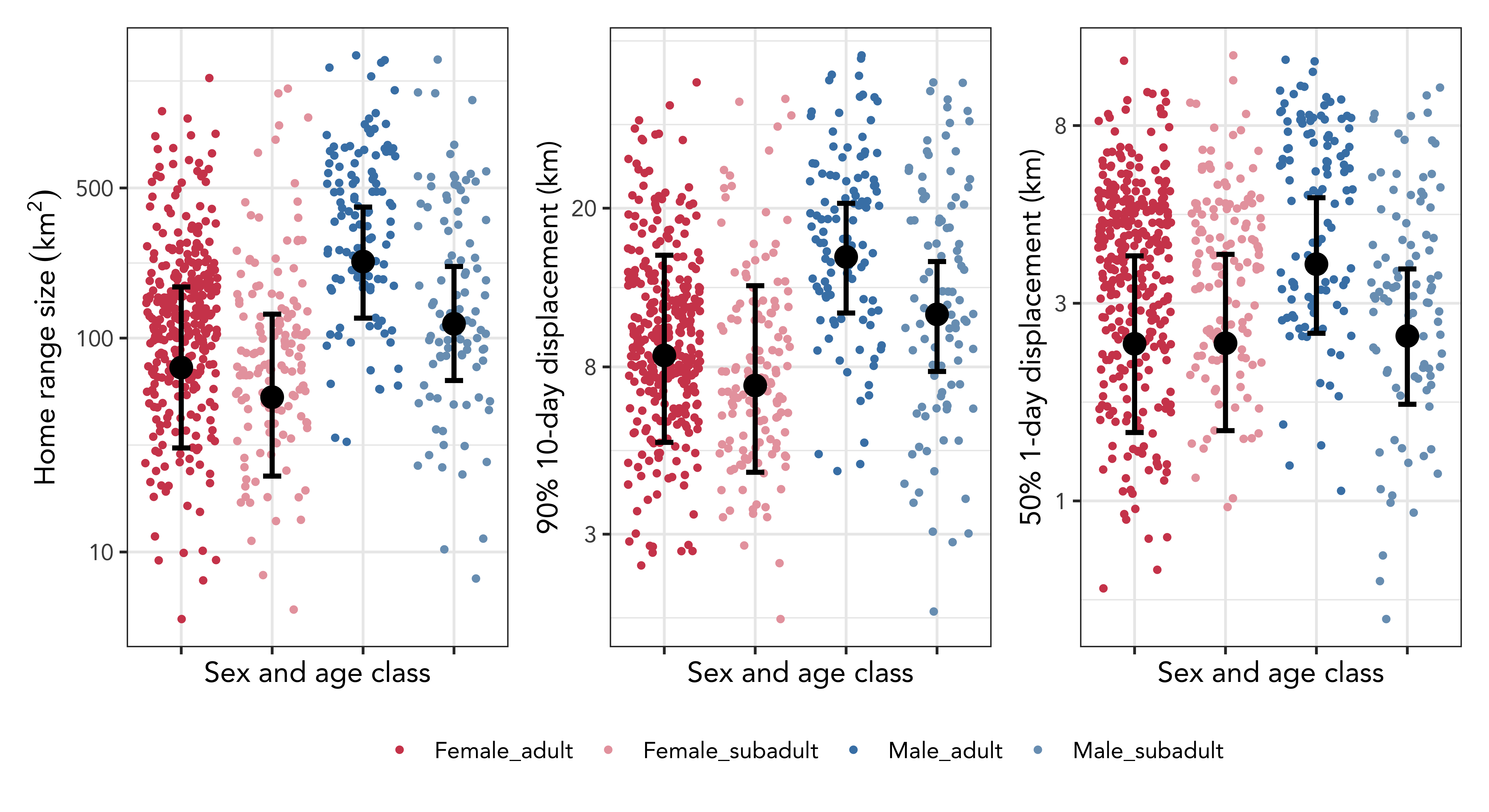

**Figure S5.** Sex and age class specific space use of bear, measured as home range size, long distance displacements, and daily routine displacement distances. We categorized bears as subadult up until age 4 and as adult when they were estimated to be 5 years or older.

As expected, age effects on movement were larger in males than in females but not in the direction predicted, i.e., we observed larger home ranges in adults that in subadults. Since we already filtered out bears that showed obvious directed dispersal, we apparently only retained subadult individuals that occupied a home range. While age effects were apparent, we refrained from including age class into the main model because a) several individuals were missing any age information (n = 71), and b) methodology to estimate age and reporting standards varied widely across our dataset.

**Appendix 5. Sex – specific movement responses**

In the main text we test how space use of brown bears was affected by landscape level covariates of food availability, climate, and human disturbance. For the brown bear, sex-specific movement responses to the environment have been shown previously, that is, female and male bears adjust movement differently to the environment or human pressures. Most of these studies show differences in landscape level habitat selection (i.e., where do males versus females establish home ranges) and fine scale habitat selection (i.e., how do males versus females utilize the habitat within these home ranges). For example, females have been shown to occupy home ranges closer to human settlements while male home ranges are more remote (Nellemann *et al.* 2007) and within their home ranges, females with cubs tend to select for human built up as a human shield against male infanticide (Rode, Farley & Robbins 2006; Steyaert *et al.* 2016). While we here do not study habitat selection, we could expect that males and females also differ in their movement responses towards their home range composition.

Specifically, bears may show sex-specific movement responses to human disturbance, with females reducing movement more strongly in areas of greater human influence as compared to males. Bears may also show sex-specific responses towards the proportion of early successional forest. While forest clearings provide abundant food, their opennes may also facilitate faster movement. We expected that male bears can traverse their home range faster when encompassing a greater proportion of early successional forest. Last bears occupied smaller home ranges and moved less in areas of higher vegetation productivity, yet differences in the diet of males and females could make a sex-specific reponse plausible.

We here explored sex-specific responses of European brown bears to human footprint, annual vegetation productivity, and the proportion of early successional forest within their home range. We refitted the main models for home range size, 10-day displacement and 1-day displacement with interactions between sex and either of the three predictor variables, respectively, resulting in 12 additional models. We found no evidence for sex-specific responses to human footprint or annual vegetation productivity (**Figure S6, S7, S8**). We found that male home range size decreased more strongly with increasing proportion of early successional forest in a bear’s home range as compared to females, which was counter our expectation (**Figure S6**). We found no sex-specific effect of the proportion of early successional forest on long-distance movements (90% 10-day displacements, **Figure S7**) or routine movements (median 1-day displacements, **Figure S8**) .

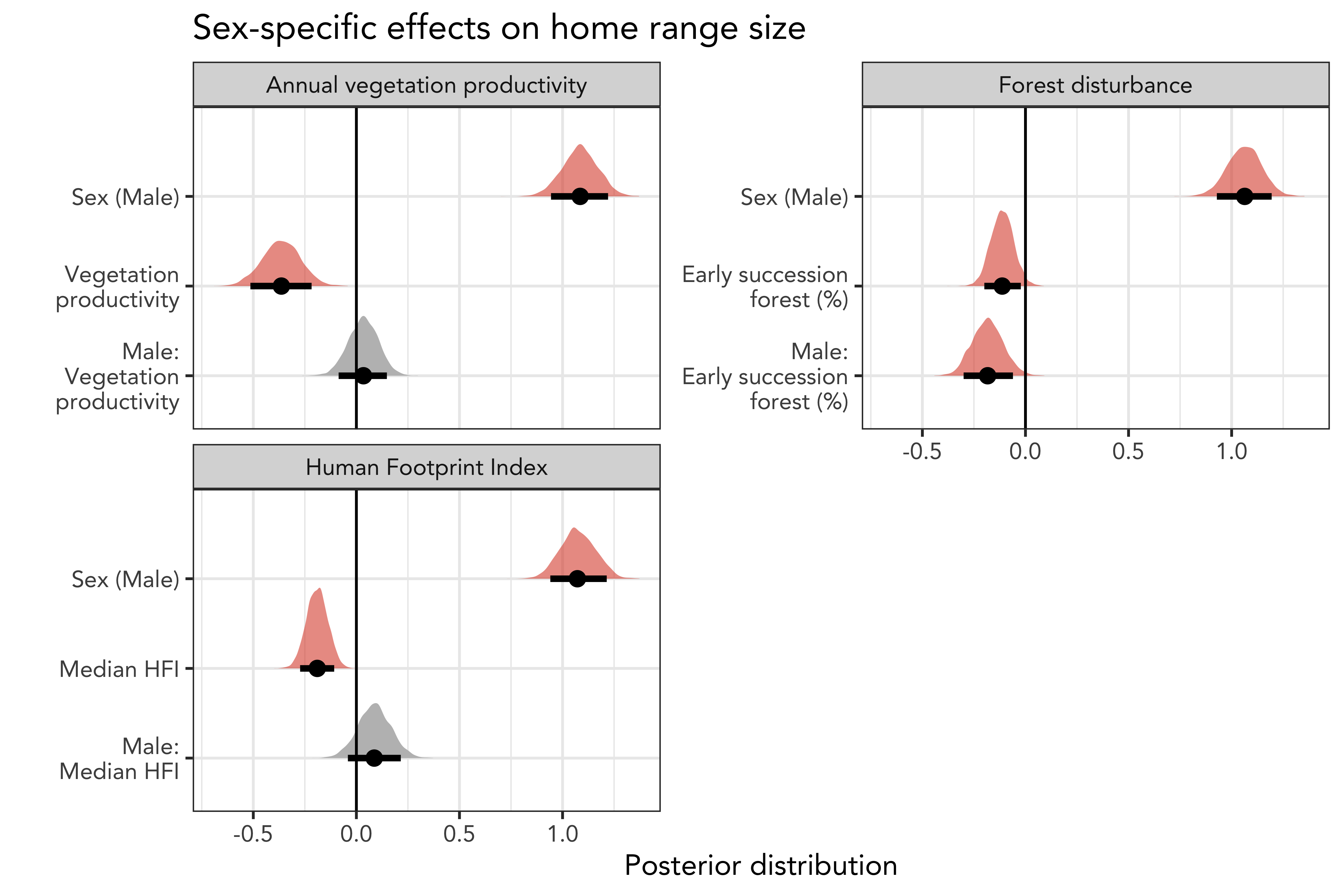

**Figure S6** Sex-specific effects of annual vegetation productivity, the proportion of early successional forest (i.e., after clearings), and human footprint on home range size. Males occupied larger home ranges than females and home ranges were smaller with increasing vegetation productivity, and human footprint, however there was no sex-specific decrease in home range size. Home ranges were also smaller with increasing proportions of early successional forest and males decreased home range sizes more strongly than females.

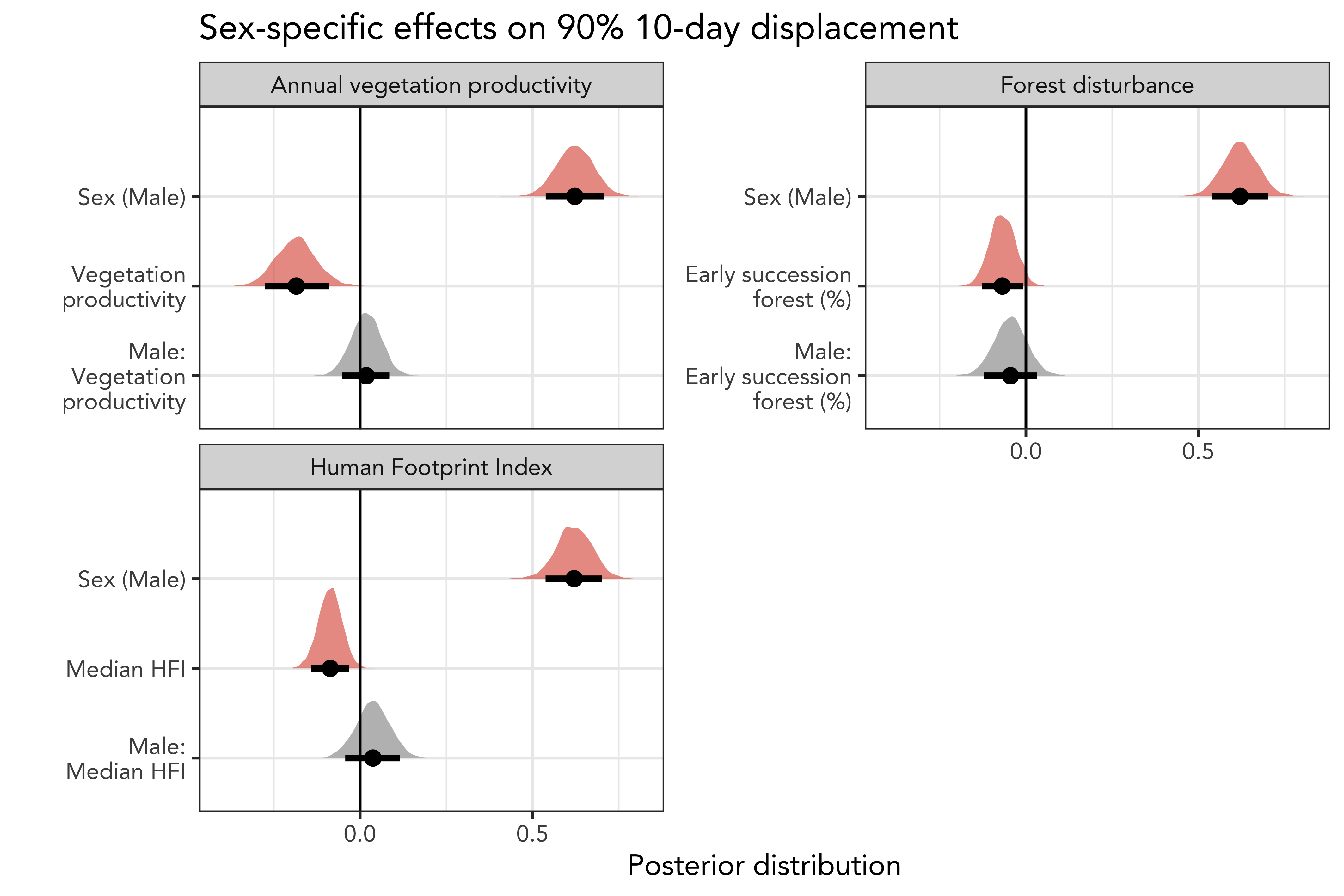

**Figure S7** Sex-specific effects on long-distance 10-day displacement distances. Males moved over longer distances than females. All bears moved less in areas of higher vegetation productivity, human footprint index, early successional forests, and terrain ruggedness. There was no sex-specific decrease in long distance movements.

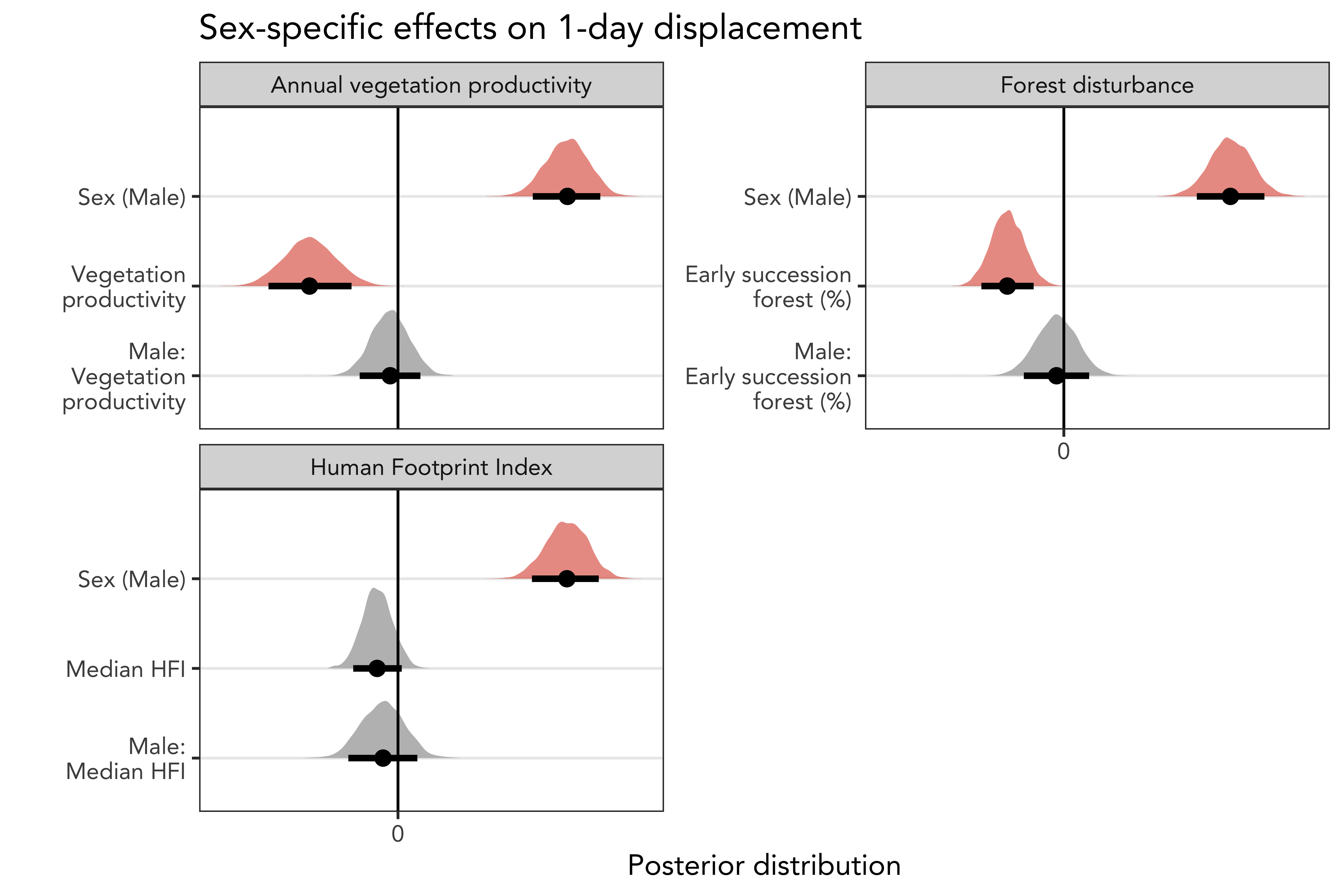

**Figure S8** Sex-specific effects on routine 1-day displacement distances. Males moved over longer daily distances than females. All bears moved less in areas of higher vegetation productivity and early successional forests, but after controlling for sex specific effects the main effect of HFI was not significant anymore. There was no sex-specific decrease in routine daily movements.
